# Frequent seasonal reassortment between high and low path viruses drives the diversification of influenza A/H5N1

**DOI:** 10.64898/2026.04.17.719307

**Authors:** Lambodhar Damodaran, Joseph A. Lewnard, Gregg S. Davis, Sara Y. Tartof, Louise H. Moncla, Nicola F. Müller

## Abstract

Since 2021, highly pathogenic (HPAI) H5N1 viruses have spread across the Americas, diversifying via reassortment into new genotypes that have spilled into humans and livestock, raising fears of a new influenza pandemic. Pandemic lineages are typically associated with reassortment, but we currently have limited understanding of where and when reassortment is expected to occur, which limits our ability to assess pandemic risks. Using a dataset of 9,052 full-genome sequences, we show that reassortment and novel genotype formation are associated with seasonal variation in low pathogenicity avian influenza (LPAI) cases and with the spatial and host distributions of viral transmission. We pinpoint ducks, geese, and the Central flyway as frequent sources of new genotypes, and show that reassortment rates vary seasonally, driven by mixing between high- and low-pathogenicity viruses. Cattle spillover genotypes (B3.13 and D1.1) evolved during periods of high reassortment, implicating reassortment as a common occurrence in lineages evolving during particular time periods. Together, these findings reframe reassortment as a predictable ecological process, with direct implications for how surveillance and pandemic risk assessment should be designed.

## Introduction

Since late 2021, highly pathogenic avian influenza (HPAI) clade 2.3.4.4b viruses have spread across the Americas, resulting in a panzootic that has caused mass die-offs among wild birds and mammals [23], extensive culling of domestic birds [12], and at least 70 human spillover infections. Recent evidence suggests that transmission of clade 2.3.4.4b viruses is increasingly driven by wild birds [16, 27, 42], leading to seasonal outbreaks and more frequent reassortment [40]. Since the incursion into North America, clade 2.3.4.4b viruses have reassorted with low-pathogenicity avian influenza (LPAI) viruses that circulate enzootically, forming new genotypes that have successively replaced prior circulating diversity [40, 43]. In 2024 and 2025, a subset of reassortment events generated novel genotypes that spilled into domestic dairy cattle (B3.13 and D1.1 genotypes) [31], spurring new waves of poultry outbreaks and associated human spillovers [15, 19], underscoring the importance of understanding the processes that drive reassortment and novel virus formation.

Reassortment enables influenza viruses to reshuffle gene segments between different ancestral lineages, allowing for the rapid formation of novel genetic constellations. Of the influenza pandemics, all but one resulted from re-assortment events. Reassortment between co-circulating seasonal influenza viral variants can lead to viruses with increased fitness that spread rapidly throughout the population [26]. Among influenza viruses circulating in animals, reassortment periodically results in novel zoonotic viruses that spill into humans, including swine-variant strains, avian H7N9, H5N6 [7, 30], and H10N8 [13], and recent reassortant clade 2.3.2.1 [39] and 2.3.4.4b H5N1 viruses. Reassortment requires co-infection with two or more lineages of different parental strains within the same host and cell. As such, reassortment rates should scale with prevalence, the frequency of infection within a population. Co-infection and prevalence may also be influenced by factors such as host-specific differences in susceptibility to viral strains, spatial and geographic heterogeneity in host movement and aggregation patterns, and variation in host immunity. However, reassortment inference typically remains largely qualitative, with current approaches focused on measuring whether two clades are separated by at least one reassortment event, precluding quantitative measurements of reassortment and direct hypothesis testing [5, 28]. As a result, the ecological and epidemiologic factors that drive reassortment remain poorly resolved, limiting accurate risk assessment and surveillance activities.

Here, we use state-of-the-art Bayesian phylodynamic inference to to trace how novel H5N1 genotypes formed across North American species and flyways. We show that while genotype formation is well-described by host-specific transmission patterns, ducks and geese collectively account for the vast majority of novel genotype generation. We identify the Central flyway as particularly important for genotype formation, consistent with its known role as a key breeding ground for North American Anseriformes. We then demonstrate that time-varying reassortment dynamics can be expressed as a function of the pathogen’s prevalence in a population and its transmission rate. Using a novel inference approach to reconstruct the time-varying nature of reassortment, we reveal extensive reassortment between co-circulating high- and low-pathogenic avian influenza virus lineages, not restricted to any specific subtype, highlighting the crucial role of low-pathogenicity avian influenza viruses in the emergence of the novel H5N1 genotypes. We find evidence for seasonally varying reassortment rates and show that the emergence of key genotypes (B3.13 and D1.1) arose following periods of high reassortment.

Overall, our results support a view of reassortment that is ubiquitous, seasonal, and driven by the hosts and geographic locations that support transmission, suggesting that surveillance could be better targeted. Our results suggest that the same force that drives spillover may also drive reassortment (prevalence), and suggest that the mere presence of a reassortment event in the recent ancestry of a viral clade itself is not necessarily evolutionarily significant.

## Results

### New HPAI influenza genotypes emerge across all North American Flyways

North American clade 2.3.4.4b viruses are classified into genotypes that represent novel introductions (denoted as “A” genotypes) or reassortants (B, C, and D genotypes) [43], making genotypes a proxy for successful reassortant viruses. To trace the spatial and host distribution of H5N1 genotype formation, we compiled all available full genome sequences from the GISAID database submitted before 2025-06-23. We assigned each sequence to the US Fish and Wildlife migratory flyway of sampling (Atlantic, Mississippi, Central, Pacific) using location metadata, and to one of 9 host groups using species-level host information. Following removal of temporal outliers, we assembled a dataset of 9,052 complete H5N1 genomes (see Materials and Methods for details), and inferred phylogenetic trees for each gene segment individually, using the ‘TargetedBeast’ package in BEAST2 [9]. TargetedBeast utilizes more efficient tree proposals, enabling fully Bayesian phylodynamic inference from these large datasets, allowing us to infer tree topologies and evolutionary parameters simultaneously. For each analysis, we jointly inferred host and migratory flyway as discrete traits across the posterior set of trees, and quantified transitions across the trees using stochastic mapping [25].

Consistent with prior reports [11, 16], HPAI H5N1 was introduced independently into the Atlantic and Pacific flyways, and then spread from the Atlantic flyway westward (Figure 1A). H5N1 cases and sequences from 2021-2025 derive from all 4 flyways (Figure 1B), highlighting the ubiquity of these viruses across the continent. Because reassortment (and by proxy, genotype formation) requires co-infection within a host organism, we hypothesized that genotypes would form in hosts and flyways proportionally to the transmission rates they support. To test this, we inferred the time to the most recent common ancestor for each segment for each genotype, and reconstructed the most likely flyway origin (Figure S8–S9). To approximate the degree of transmission occurring across flyways, we mapped the proportion of all tree edges in each flyway in rolling, two-week time windows, and compared the proportion of tree edges (a proxy for transmission) to the proportion of originating genotype segments (a proxy for reassortment) from each flyway; the observed/expected ratio corrects for potential biases from sampling density and coalescent structure (see Methods for details).

**Figure 1.**
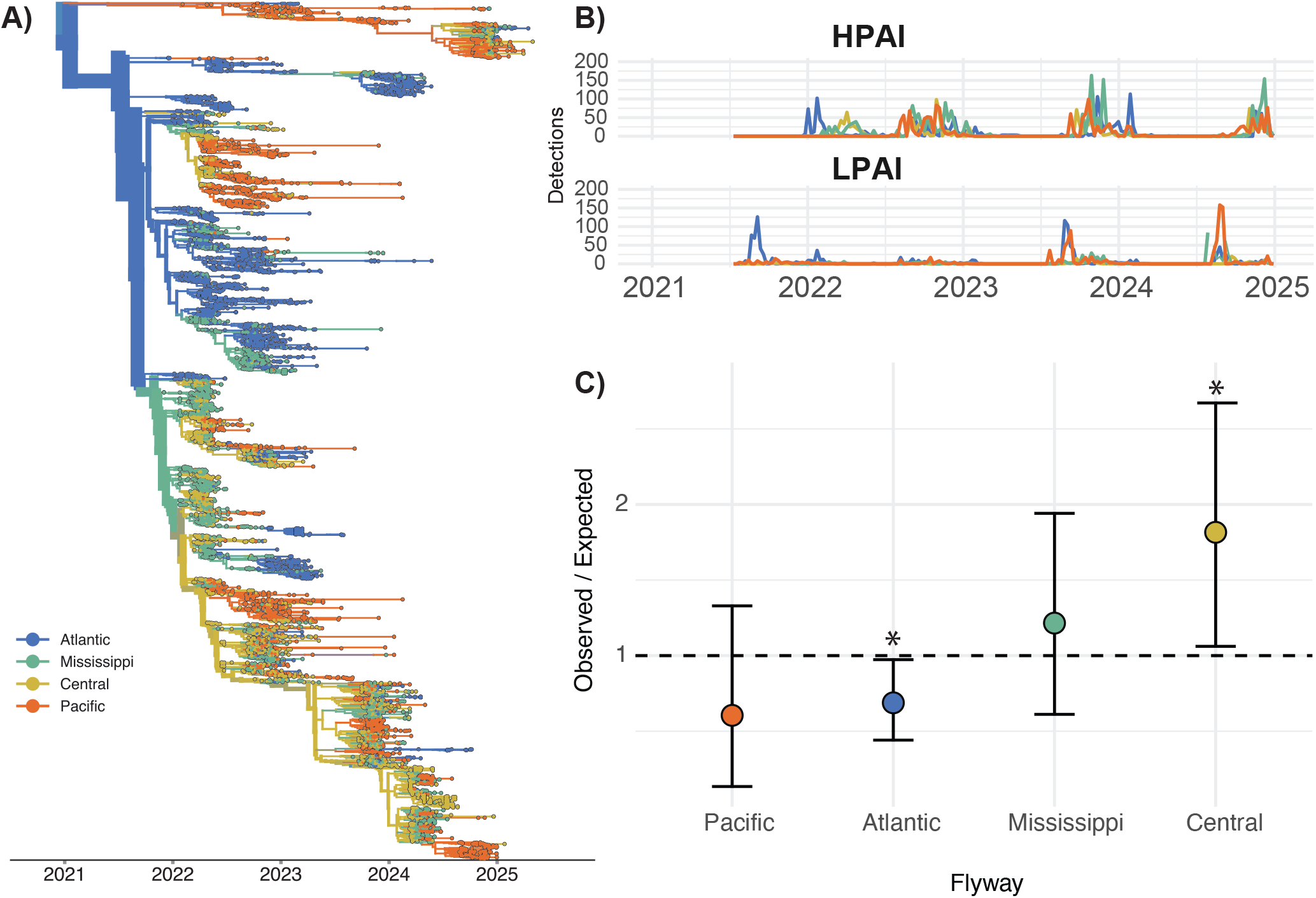
HPAI H5N1 clade 2.3.4.4b spread widely across North America. **A** Here we show the maximum clade credibility tree for 9052 HA sequences of HPAI H5N1 colored by inferred migratory flyway. We inferred the flyways using an asymmetric discrete trait diffusion model. Trees for all other segments are shown in Figure S1–S7. **B** Detections of HPAI and LPAI in North America, colored by migratory flyway, published by USDA Animal & Plant Health Inspection Service (APHIS). **C** Results of the observed/expected ratio for genotype emergence from a given migratory flyway. The 95% confidence interval from bootstrap resampling of genotype are indicated by error bars. We indicate values that were statistically significant (p¡0.05) with a ∗.

During the first year of the panzootic (2021-2022), genotype formation approximately tracked the westward movement across migratory flyways, with close concordance between tree edges and genotype segment origins (Figure 1B, Figure S1–S7). Following mid-2022, genotypes increasingly derived from the Central flyway, deviating from underlying expectations based on transmission rates alone (Figure 1C, Figure S8–S10). Genotype segments originated from the Central flyway 1.82 times more frequently than expected (95%CI = 1.06-2.5, p-value = 3.8E-02) (Figure S11). Compared to the expected origin of genotype segments, segments originated from the Atlantic (observed/expected = 0.69, range=0.44-0.97) less frequently than expected, and formed in the Mississippi (observed/expected = 1.21, range=0.61-1.94) and Pacific (observed/expected = 0.60, range=0.13-1.33) at approximately expected frequencies (Figure S11). Thus, while transmission and genotype formation occurred across all 4 flyways, genotype segments originated most frequently from the Central flyway.

### Genotype formation is proportional to host-specific transmission, but enriched in geese

Past work has identified Anseriforme and Charadriiforme species as key drivers of avian influenza virus evolution [18] [22].To determine the contribution of host species to viral transmission and genotype formation, we next classified sequences into 9 host groups and inferred phylogenetic trees with host annotated as a discrete trait. Repeating the approach from above, we inferred the most likely host source for each genotype segment and compared transmission to genotype formation over time.

Phylogenetic trees across all segments support a key role for ducks in long-term and persistent transmission of HPAI (Figure 2A, Figure S12–S18). Ducks were the most commonly inferred host across the tree, particularly along the tree backbone and deep internal nodes, supporting an important role for sustained transmission in these species. Enumeration of yearly transmission patterns show that transmission among ducks frequently occurs earlier than transmission within other species, preceding yearly transmission peaks in geese, galliformes, and other species (Figure 2C). These results are compatible with a model in which ducks, potentially via early season migration, play a role in early season movement and dissemination of viral lineages, a hypothesis which could be tested through active surveillance [3, 14, 32].

**Figure 2.**
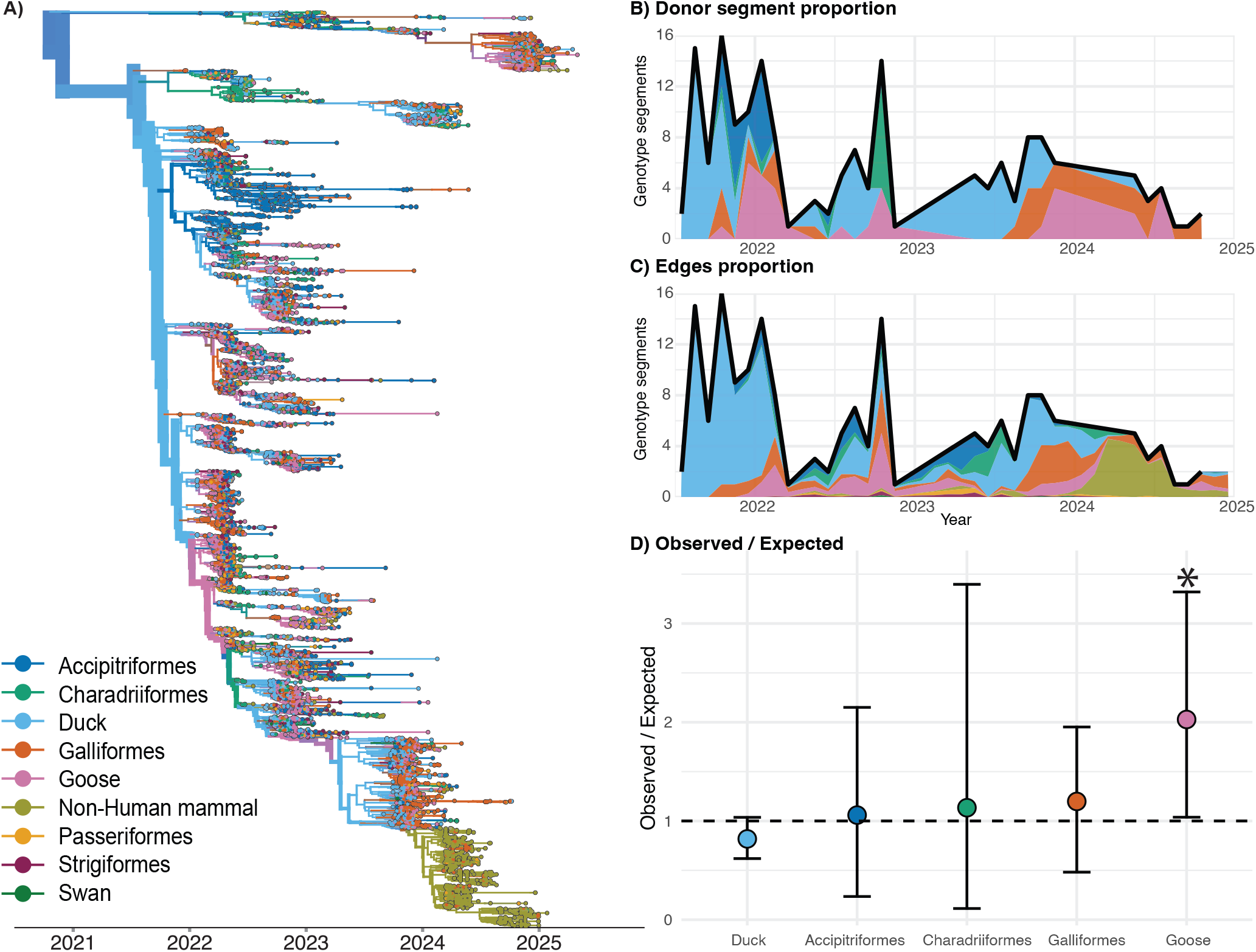
Host dynamics of HPAI H5N1 clade 2.3.4.4b in North America. **A** Maximum clade credibility tree for 9052 HA sequences of HPAI H5N1 colored by inferred host. Trees for all other segments are shown in Figure S12–S18. **B** Number of genotype segments in monthly time-bins with the proportion of segments inferred from a given host colored below the line. **C** Number of genotype segments in monthly time-bins with the proportion of edges across the phylogeny coming from a given host colored below the line. Notably, while the proportion of inferred genotype segments from galliformes increased in 2024, this likely reflects the limited data from wild birds in this time period (Figure S19). **D** Observed/Expected ratio for genotype emergence from a given host. The 95% confidence interval from bootstrap resampling of genotype are indicated by error bars. Values with statistical significance are starred.

Among the 20 reassortant (non-A) genotypes we analyzed, 18 included gene segments that were inferred to derive from multiple host groups (Figure S20, S21), suggesting frequent viral mixing between species. However, genotypes formed across most species at frequencies that are expected from their transmission rates (Figure 2D). While some segments were inferred to derive from galliformes, raptors, and shorebirds, these were transient and rare, supporting a more limited role in genotype formation that can be explained by lower transmission rates (Figure 2B). Similarly, we find no genotype segments originating in songbirds, owls, and swans, consistent with expectations from transmission rates alone, and supporting a limited role for these species in genotype formation.

Anseriformes were inferred as the most frequent sources for viral gene segments by far, with at least one segment deriving from ducks or geese across all analyzed genotypes. Ducks accounted for 46% of all genotype segments—the largest share by far from any host species—supporting an important role for genotype formation that can be explained by high transmission rates. The two exceptions to these patterns were geese and nonhuman mammals. Geese supported lower transmission overall, but contributed twice as many genotype segments as expected (O/E = 2.03, 95% CI= 1.04-3.32, p= 4.2E-02). We find no evidence for genotype segments originating from nonhuman mammals, despite the high degree of transmission in dairy cattle in 2024 (observed = 0, expected = 6.9%), confirming a limited role for dairy cattle in generating novel genotypes to date (Figure S22). Taken together, these data support a strong link between genotype formation and viral transmission patterns among species, with Anseriformes inferred as key sources of new viral genotypes. Our data also support geese as potential outsized contributors to reassorted segments, though additional work is necessary to confirm these results more broadly.

### High reassortment rates in HPAI H5N1 vary seasonally with LPAI cases

Reassortment is intrinsically linked to co-infection, and is only detectable if it occurs between viral lineages that are genetically distinct. We next derived the relationship between infection prevalence and reassortment rates using a compartmental (susceptible-infectious-recovered) model that allows for co-infection by multiple viruses in the same host (Figure S23). In this model, an infected individual (I) can infect a susceptible individual (S) with a rate of 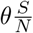 denotes the transmission rate and 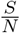 represents the proportion of the population susceptible to infection. Co-infection events then happen at a rate 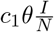 describes the proportion of encounters between two infected individuals that lead to a co-infection event. Upon co-infection, a reassortant virus is created in *c*_2_ of all cases. We show in *Methods and Materials* that we can express the time-varying reassortment rate in a coalescent with reassortment model as a function of the transmission rate, c, and the disease prevalence 1.

We next expanded the Coalescent with Reassortment (CoalRe), implemented in BEAST2, to infer time-varying reassortment rates. To enable direct interrogation of how epidemiological and ecological factors influence reassortment rates, we further implemented the ability to infer predictors of the time-varying reassortment rate. We downloaded complete whole-genome HPAI H5Nx sequences from viruses sampled in North America submitted before 2025-06-23 from the GISAID database, and generated randomly sub-sampled datasets of 350 HPAI samples for computational tractability. We then reconstructed the reassortment dynamics of North American H5Nx viruses circulating between 2021 and 2025.

As shown in Fig 3A, reassortment occurs frequently across the history of HPAI in North America. Reassortment rates vary over time, with an initial peak in reassortment in fall of 2021, around the time when the 2.3.4.4b clade was first introduced into North America [11]. Because inferred reassortment networks include the histories of all gene segments, these events may also include reassortment events in LPAI lineages that later donated segments to HPAI lineages. Reassortment rates were lower in 2022, before peaking again in the summers of 2023 and 2024 (see Fig 3E) at a rate of about 1 event per lineage per year. These summer peaks in reassortment precede fall surges in H5N1 cases, and coincide roughly with times of high LPAI case detections. We find modest support for detected LPAI cases, but not HPAI cases, as predictors for reassortment rates(see Fig 3B), though case detections alone cannot fully explain the observed dynamics. These data suggest that reassortment rates vary seasonally, and are best, but not completely, predicted by time-varying LPAI cases.

**Figure 3.**
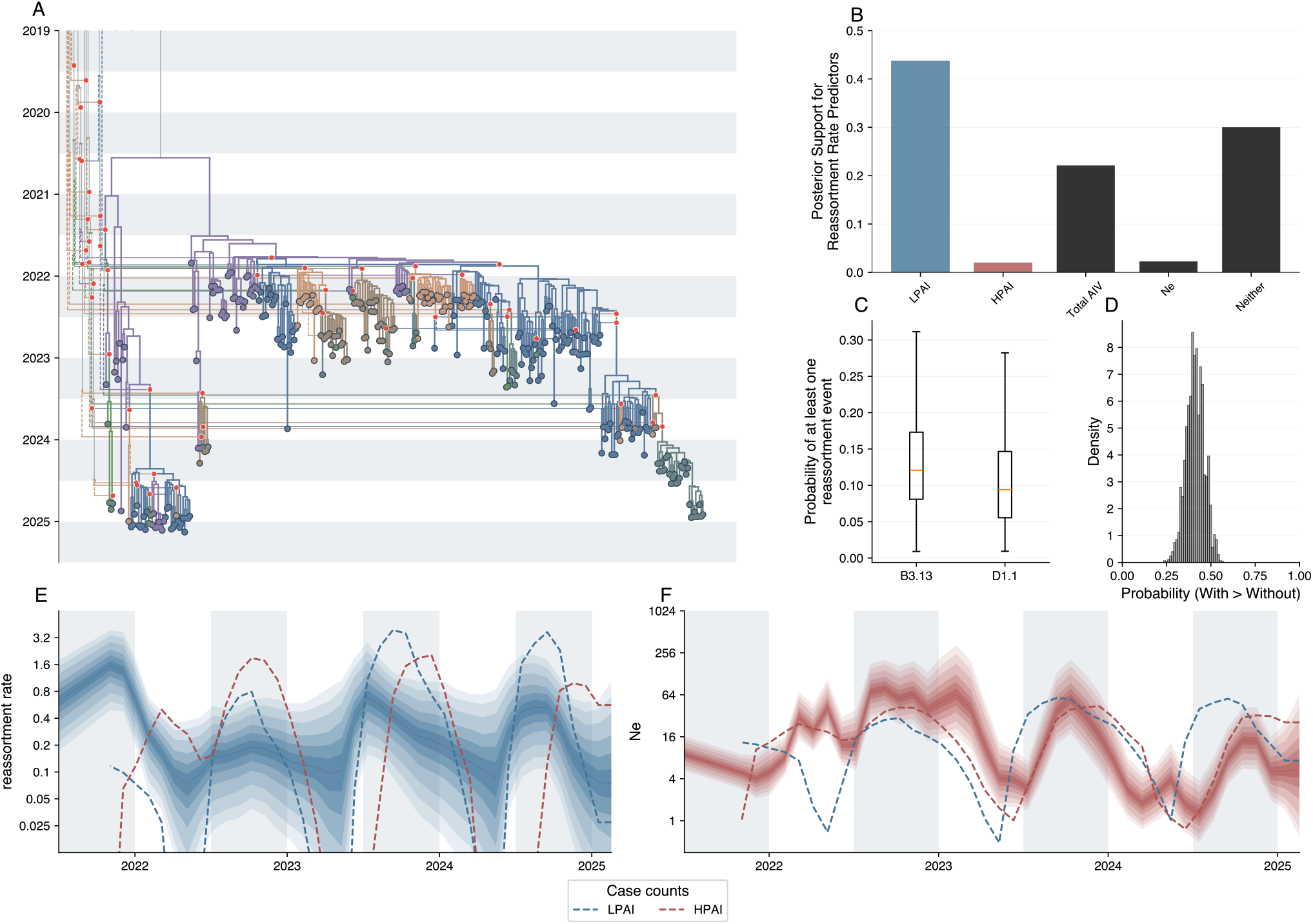
Reassortment rates in HPAI Influenza A/H5N1 in North America. **A** Maximum clade credibility network reconstructed from influenza A/H5N1 sequences collected in North America. The network is organized by HA segment. Each red dot denotes a reassortment event. **B** Posterior support for cases predicting reassortment rates. **C** Probability that there was at least one reassortment event in the half a year preceding the most recent common ancestor of the B3.13 and D1.1 sequences in the datasets computed using the inferred reassortment rates. **D** Probability distribution of lineages with reassortment events having more offspring than their sibling lineages without. **E** Time varying reassortment rates with the different confidence intervals highlighted by the different shadings. **F** Time varying effective population sizes with the confidence intervals highlighted by the different shades.

If reassortment rates are determined purely by prevalence, then reassortment and prevalence should peak at the same time. To explore mechanistic explanations for the observed offset between reassortment rates and HPAI and LPAI cases, we next investigated scenarios where the prevalence does not directly correspond to time-varying reassortment rates. We simulated networks under SIR and SIS (susceptible infectious susceptible) models to mimic outbreaks under different epidemiologic scenarios. Under SIR dynamics, outbreaks are limited by the pool of susceptible individuals, resulting in a reduction in transmission due to a loss of susceptible hosts. In contrast, SIS models mimic a scenario in which an outbreak is limited instead by reductions in transmission rates, which could arise due to seasonal changes in contact rates or aggregation among individuals. Said another way, the SIS model mimics a scenario in which the underlying transmission rate is reduced, while the SIR models mimic a scenario in which the effective transmission rate (i.e., how many infections will find hosts) is changed. To investigate whether reassortment dynamics varied between these scenarios, we simulated networks under 3 SIR models: SIR with co-infection; SIR with co-infection and superspreading; and SIR with population structure and superspreading; and an SIS model with time-varying transmission rates (Figure S24, Figure S25). We find that while reassortment rates under SIR dynamics result in reassortment rates that match trends in prevalence, that SIS dynamics (limiting outbreaks by reducing transmission) result in reassortment rates that can be markedly lower, and that peak earlier than prevalence. Together, these results suggest that ecological and epidemiologic factors (e.g., host movement, aggregation, immunity) may augment the timing and intensity of reassortment.

### Reassortment was expected preceding recent, emergent genotypes

Reassortment of H5Nx viruses has led to the constant emergence of new viruses that occasionally cause new waves of spillovers to humans and agriculture. Notably, the B3.13 and D1.1 lineages emerged following reassortment and spilled into dairy cattle and humans, leading to speculation that reassortment itself may have provided a fitness benefit. Given the high rates of reassortment we observe, we next asked how probable it is to observe a reassortment event preceding a lineage that sweeps the population, even if the reassortment event provided no fitness benefit. We find that the expected probability of a reassortment event in the 0.5 years preceding emergence of the B3.13 and D1.1 lineages, under the null hypothesis of no fitness benefit due to reassortment, is 15%, solely based on the reassortment rates during those times (see Fig 3C). These data imply that reassortment is highly probable in the evolutionary history of viruses evolving during particular time periods, even if it provides no fitness benefit. Thus, reassortment in new lineages may not necessarily indicate surprising events, but rather expected outcomes of viral circulation during particular times and places.

### Repeated reassortment between HPAI and LPAI drives diversification

The correlation between HPAI reassortment and LPAI cases implies that LPAI viruses may act as important donors to HPAI reassortants. We next explicitly included HxNx LPAI isolates from North America into our analyses, and reconstructed the joint evolutionary history of HPAI and LPAI lineages from all eight segments. Adding the LPAI isolates allowed us to classify each network lineage in the posterior distribution as either HPAI or LPAI based on the hemagglutinin segment to which the network lineage corresponds. We then mapped each time a reassortment event introduced a segment from an LPAI lineage into an HPAI lineage to formally estimate the relative reassortment rates among HPAI and LPAI lineages. Figure 4A shows the hemagglutinin tree together with the timing of LPAI segment introductions into HPAI lineages; segment-specific trees for all non-HA segments are shown in Figure S26.

**Figure 4.**
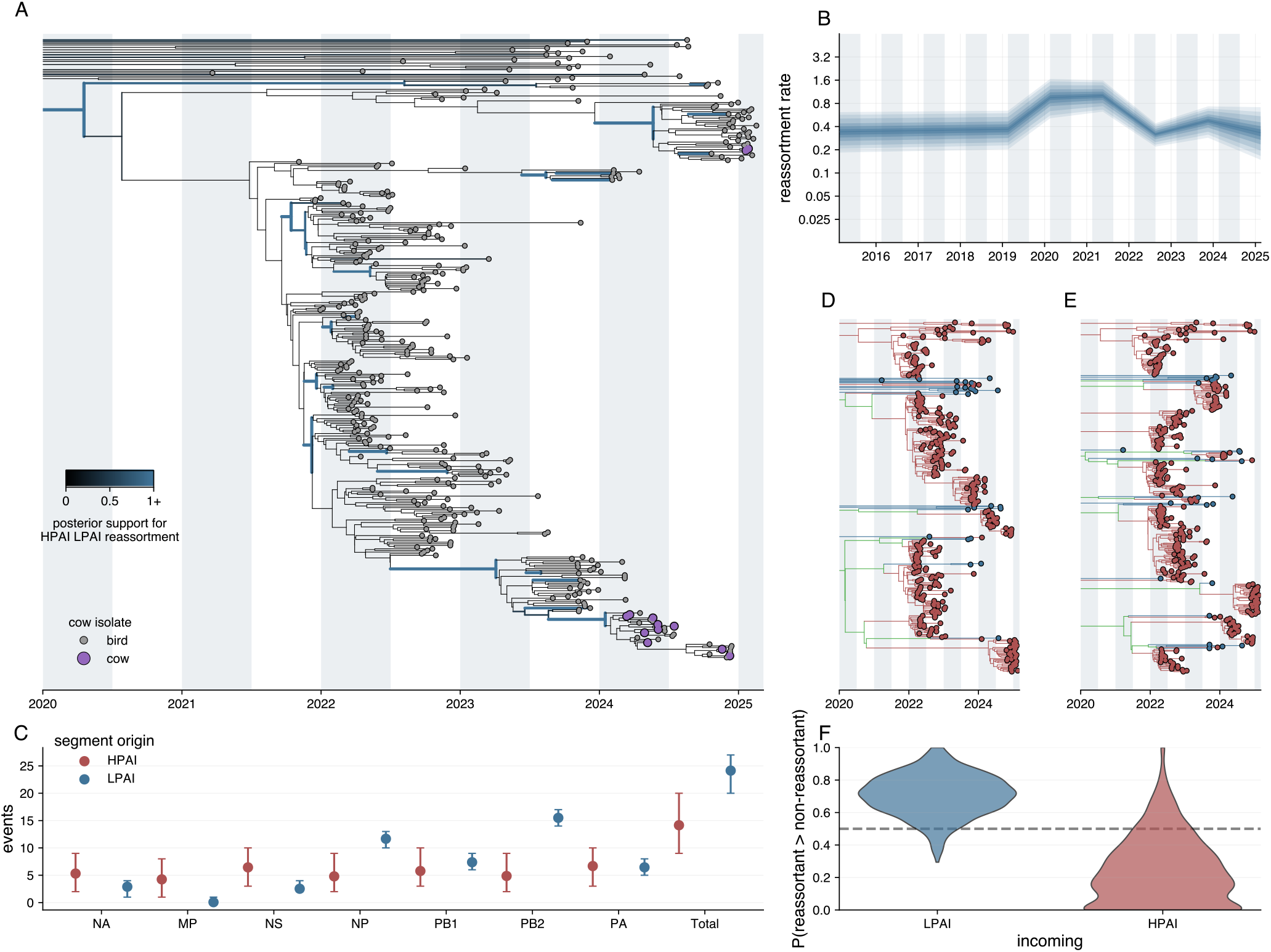
Analysis of reassortment of low pathogen lineages with high pathogenic influenza A/H5N1 lineages in North America. **A** Maximum clade credibility tree of the hemagglutinin (HA) segment. On the branch, we map the posterior number of reassortment events between HPAI and LPAI lineages, shown by the green color. Tip colors indicate cow isolates (purple) and bird isolates (gray). **B** Temporal dynamics of reassortment rates estimated using skygrowth analysis, showing median rates (thick line) and confidence intervals (shaded ribbons). **C** Co-reassortment events by segment, showing the number of events where segments originated from HPAI (red) or LPAI (green) lineages, with error bars representing quantiles. (D) Maximum clade credibility tree of the nucleoprotein (NP) segment, colored by lineage type (HPAI: red, LPAI: green, unknown: green). **E** Maximum clade credibility tree of the polymerase basic protein 2 (PB2) segment, colored by lineage type. All trees are time-scaled with alternating shaded regions indicating 6-month intervals. **F** Comparison of the number of offspring between sibling edges, one with a reassortment event, the other without one. For each iteration in the posterior, we bootstrap the events used for the calculation to avoid being biased by single events.

We find evidence for frequent reassortment between HPAI and LPAI lineages, aligning with above results. Enumeration of the posterior number of segments donated to HPAI lineages showed that reassortment events involving NA, MP, and NS were rare, consistent with strong linkage between these segments and HPAI H5 genes (Figure 4C). NP and PB2 were the most frequently reassorting segments, with multiple clades of NP and PB2 gene from HPAI lineages nested within the diversity of low pathogenic lineages (Figure 4D&E ). Thus, HPAI reassortment in North America is overwhelmingly driven by acquisition of novel LPAI genes, with PB2 and NP acquisitions occurring with the greatest frequency. One important caveat is that HPAI lineages are less diverse than LPAI lineages, leading to a potential detection bias that would favor detecting reassortments involving LPAI viruses. Underestimation of HPAI-HPAI reasssortment events relative to HPAI-LPAI events could inflate the apparent role of LPAI as a predictor of reassortment rates. Given the high apparent propensity for HPAI-LPAI reassortment events, we next investigated whether HPAI-LPAI reassortment events were associated with a fitness benefit. We quantified the number of edges that descended from reassortant and non-reassortant branches as a measure of fitness, with the expectation that more fit edges will produce more offspring. We then classified each reassortment event into whether the incoming segments originated from an HPAI or an LPAI virus. As shown in Figure 4F, reassortment events in which the incoming segment originates from an LPAI lineage tend to have more offspring than their siblings, which did not reassort, while events where the segments originate from HPAI lineages tend to have fewer. These results are compatible with a model in which HPAI-LPAI reassortment events are associated with fitter viruses that disseminate to more offspring. However, this effect can arise both if reassortment between HPAI and LPAI is innately beneficial or if HPAI-LPAI reassortment is simply correlated with other features that underlie fitness (e.g., circulation in particular hosts or environments). Regardless, these results suggest HPAI-LPAI reassortment as a potential feature of fit HPAI lineages in North America.

## Discussion

Reassortment is often seen as a signal of pandemic risk, yet the processes governing when, where, and in which hosts it occurs remain poorly understood. Here, we show that reassortment of HPAI H5N1 in North America is frequent, seasonal, and largely predictable from ecological and epidemiological factors. We infer that these re-assortment dynamics are driven by LPAI co-circulation and are concentrated in specific hosts and migratory flyways. These findings suggest that reassortment is not a surprising evolutionary event, but an expected outcome of high viral circulation during particular times and places, with direct implications for how surveillance programs and pandemic risk assessments should be designed. The establishment of HPAI H5N1 clade 2.3.4.4b in North American wild birds, and its subsequent spillover into dairy cattle and humans, has intensified concern that reassortment events themselves signal dangerous evolutionary transitions. Our data allow us to evaluate this assumption directly.

Reassortment has historically been described qualitatively, with few approaches capable of linking it mechanistically to ecological or epidemiological hypotheses. By pairing a large, representative dataset of 9,052 H5N1 genomes with new phylodynamic methods enabling fully Bayesian inference at this scale, we can reconstruct how genotype formation connects to transmission across species and flyways. Using a mechanistic SIR model incorporating co-infection dynamics, we show that reassortment rates scale with prevalence, providing a direct theoretical link between epidemiological dynamics and evolutionary outcomes. We then incorporate these insights into Bayesian phylodynamic inference to reconstruct time-varying reassortment dynamics. We find that genotype formation and reassortment dynamics are partially predicted by seasonal LPAI case counts and by the spatial and host distribution of viral lineages. Reassortment rates peak in summer, preceding fall surges in HPAI cases, and predominantly involve acquisition of novel LPAI gene segments by HPAI lineages. Importantly, LPAI cases, rather than HPAI cases, show modest support as the strongest predictor of reassortment rates, implicating LPAI circulation as an important driver of HPAI diversification. We specifically investigate these by looking at the emergence of two reassortant clades that spilled over into humans. The probability of observing a reassortment event in the emergence of B3.13 and D1.1 was 15% under the null hypothesis of no fitness benefit from reassortment — meaning that reassortment in the ancestry of these lineages is compatible with random chance given the small sample size (n=2), though both lineages having reassortment is slightly above the expected 0.3 under the null. Together, these results support prevalence as the key predictor of reassortment frequency, and suggest that the presence of reassortment in a lineage’s recent history is not itself evidence of unusual evolutionary pressure.

Notably, our offspring analysis (Figure 4F) shows that reassortment events involving LPAI-derived segments tend to have more offspring than their siblings, while HPAI-derived segments tend to have fewer, suggesting that the acquisition of an LPAI segment (mainly NP and PB2) is associated with a fitness benefit. This suggests that reassortment is frequent enough that its presence in successful lineages does not imply selection, yet when it does occur, LPAI segment acquisition may frequently be associated with fitness advantages. Most HPAI genotypes that have dominated in the Americas are HPAI-LPAI reassortants, supporting this conclusion. The long-term co-evolution of LPAI viruses with North American birds could mean that LPAI genes are better adapted for circulation in these hosts, thus providing a fitness benefit during reassortment. Alternatively, these effects could arise from epidemiologic or ecological features. If circulation in particular hosts or locations enhances spread, then co-circulation of HPAI and LPAI within the same high-transmission species could lead to HPAI-LPAI reassortment that correlates with, but does not cause, spread to more offspring. Regardless, our results suggest that HPAI-LPAI reassortment is associated with higher fitness viruses, though future work is necessary to disentangle the cause.

Our data support ducks as important, early-season drivers of H5Nx transmission and reassortment. These results could be impacted by the timing of surveillance efforts in different species, which could be augmented to capture more representative samples of viral diversity. Given the propensity for certain duck species, such as mallards and blue-winged teals, to migrate earlier (late summer/early fall) than other non-duck species and their lack of clinical symptoms, these data are compatible with a model in which ducks play a key role in early-season viral movement and spread [10, 14, 36]. Ducks also contributed nearly half of all segments to new genotypes, in line with the high degree of transmission they support, while geese contributed more reassortant segments than expected. These data underscore Anseriforme species as key to viral spread and mixing, but highlight that these host groups may play distinct roles in HPAI evolution. Targeted surveillance that seeks to optimize sampling based on host-specific transmission rates, reassortment propensity, and viral importation risk could be immensely useful for future viral tracking and should be explored.

Our results pinpoint ducks, geese, and the Central flyway as key contributors to the formation of novel genotypes. The Central flyway is home to the Prairie Pothole region, a heavily water-covered region that serves as the breeding ground for 60 % of all North American waterfowl and has been previously implicated in HPAI mixing [4, 40]. This region is well known among bird-watching communities, with May and June supporting the highest density of breeding duck species. We propose a model in which Anseriformes congregate in the Prairie Pothole (Central flyway) in late spring, promoting mixing with enzootic LPAI viruses and a burst of reassortment that peaks in the summer. Though the permutation test we employ assumes that genotype formation is proportional to transmission (edge counts) in the null expectation, co-infection opportunities could instead scale with aggregation density, which could explain Central flyway enrichment beyond transmission alone. Though speculative, this hypothesis unifies the results we observe across hosts, flyways, and seasons, and aligns with the known importance of this region for LPAI circulation and wild Anseriforme ecology [6, 22]. Should additional surveillance and modeling confirm this hypothesis, then enhanced surveillance in this region could potentially improve the detection of reassortant viruses before they emerge during fall surges.

Reassortment events frequently pose public health challenges because they are surprising and unpredictable. Our data show that reassortment events are expected in the recent history of two emergent lineages, B3.13 and D1.1, regardless of whether they meaningfully contribute to the fitness of these lineages. These data suggest reassortment as a frequent and ongoing process that occurs readily within the species and locations that support infection. While reassortment prediction is not yet possible, our findings suggest that clarifying the underlying ecology of viral infection could yield useful insights into the formation of reassortment viruses, potentially enabling better anticipation of the locations, times of year, and hosts where novel viruses may emerge. Central to this reframing are LPAI viruses, which our data identify as the primary source of novel segments acquired by HPAI lineages and the strongest ecological predictor of reassortment rates. LPAI viruses are currently understudied and under-surveilled relative to their apparent importance as drivers of HPAI evolution. Closing this gap through systematic LPAI surveillance may be crucial to anticipating HPAI emergence in addition to HPAI monitoring itself. As HPAI H5N1 continues to establish and evolve within North American wildlife, the ability to predict when and where novel reassortant viruses are most likely to form will be increasingly important for designing effective surveillance systems, informing vaccine strain selection, and ultimately reducing the window between novel virus emergence and public health response.

## Methods and Materials

### Data Collection

Detection data for HPAI in wild birds, domestic birds, non-human mammals, and cattle were downloaded from USDA APHIS (Access date: 2025-06-16) [1]. These detections represent laboratory-confirmed cases of HPAI, providing a date and place of specimen collection. LPAI case detection data in wild birds was downloaded from the Wild Bird Avian Influenza Surveillance Dashboard published by USDA APHIS. This data provided characterization of whether the virus tested positive as subtype H5 or H7, date of sample collection, and location of collection at the state level. Case data were summarized by date and epiweek (CDC weekly reporting standard) with the floor date in yyyy-mm-dd format (beginning of the week starting on Sunday).

We downloaded all available sequence data and corresponding metadata from the GISAID database for all North American HPAI H5Nx (clade 2.3.4.4b) viruses collected submitted before 2025-06-23 and all North American H5Nx LPAI collected between 2021-01-01 and 2025-06-23 (Date of access 2025-06-23) [38]. Only sequences with all 8 genomic segments were retained. We aligned each segment dataset using MAFFT v7.2 and visually inspected alignments using Aliview 1.81 to remove sequences causing gap regions and removed nucleotides before the start codon and after the stop codon [21, 24].

### Viral sequence data classification and phylogenetic analysis

We first aimed to determine the timing of emergence of major genotypes of HPAI in North America. We filtered all HPAI H5Nx strains by those with complete collection dates (year, month, and day), and geographic location of collection (at least state level), and classified each into subtypes using the USDA genotyping tool GenoFLU [43]. We removed temporal outliers by first estimating maximum likelihood trees for each segment using FastTree v2.1.11 and checked for temporal outliers using root-to-tip regression with TreeTime v0.11.2 [33, 37]. If a sequence was flagged as a temporal outlier in a given segment, it was removed for all other segments. This resulted in a dataset of 9,052 sequences for each genomic segment. Using the state/province level data on sample collection, each strain was classified into a US Fish and Wildlife Service migratory flyway (Atlantic, Mississippi, Central, or Pacific)[2]. This resulted in a dataset with the following number of sequences by flyway: Atlantic flyway (2447), Mississippi flyway (2352), Central flyway (1976), and Pacific flyway (2276).

We next inferred phylogenetic trees to reconstruct phylogenies and migratory flyway information simultaneously. To facilitate the large number of taxa and more efficiently search phylogenetic tree space, we used the Targeted-Beast package v1.2. and a coupled MCMC framework with hot and cold chains in BEAST v2.7.7 [8]. For each segment, we used a constant coalescent demographic model and an HKY nucleotide substitution model with gamma-distributed rate variation across sites, as well as a proportion of invariant sites with uniform prior between 0 and 1. We ran 4 independent replicates using 4 coupled MCMC chains with a chain length of 500 million states, sampling every 10,000 states [29]. We assessed convergence (ESS ¿ 200) using Tracer v1.7.2 combining the three runs with highest posterior support [35]. To infer migratory flyways onto the posterior set of trees, we modeled flyways as discrete traits using an asymmetric discrete trait diffusion model. The maximum clade credibility trees of these analyses were estimated using a posterior sample of 1000 trees using TreeAnnotator v2.7.8. To determine the migratory flyway and time of emergence of each genotype, we calculated the MRCA for each genotype and segment, and recorded the posterior probability of the ancestral state annotated at the MRCA.

Additionally, we classified sequences based on the host from which the sequence was isolated into their respective host taxonomic orders. Previous analysis has shown the importance of Anseriformes as a critical source population for virus transmission. To interrogate whether ducks and geese play distinct roles in genotype formation given their different ecologies and migration timing, we separated the Anseriformes group into two subclasses, Anseriformes-Duck and Anseriformes-Goose. The classification of strains by host order resulted in a dataset with the following numbers of sequences for each group: Accipitriformes (1477), Anseriformes-duck (1442), Anseriformes-goose (1404), Anseriforme-Swan(46), Charadriiformes (695), Galliformes (1957), Nonhuman-mammal (1054), Passeriformes(315), Strigiformes (408), and a final group of sequences annotated as ambiguous due to lack of host information (254).These host classifications were used to perform the discrete trait analysis described previously for migratory flyways.

### Permutation analysis of genotype enrichment

Because reassortment requires a co-infection, we hypothesized that spatial and host specific patterns of genotype formation would reflect spatial and host specific patterns of viral transmission. To approximate the degree of transmission within each host and flyway, we enumerated the proportion of all tree edges inferred within each flyway and host group across the MCC tree in rolling, 1-month windows. The tree edge proportions serve as a proxy for transmission, yet these edge counts may be influenced by sampling density and coalescent structure. To account for these potential biases, we compared the observed genotype segment origins to those expected values under a null model in which segments derive from flyways or hosts proportional to edges. The observed/expected ratio thus aims to correct for such biases by normalizing to transmission-weighted expectations. For each genotype segment inferred to originate within each monthly window, we inferred its flyway and host origin. For each monthly window, we then calculated the proportion of all originating genotype segments that were derived from each flyway and host. To enable direct visual comparison between the proportion of tree edges and genotype segments, we scaled both plots to the number of genotype segments originating in each monthly window.

Under a null model, in which the host and location origin of genotype formation is perfectly described by viral transmission patterns in those hosts and locations, we would expect segments for new genotypes to originate from flyways and host orders proportional to the number of inferred tree edges in those flyways and hosts. To test this hypothesis, we designed a permutation test.

For each genotype segment we defined a two-week interval around the segment’s MRCA and extracted all edges from the phylogenetic tree within this interval. For each interval, we first computed the observed number of segments of new genotypes originating from each host or flyway. To generate the expected distribution under the null, we randomly sampled one edge from each segment MRCA interval according to the relative frequency of edges in that interval. This was repeated 100,000 times. Observed counts were compared to the expectation to calculate the observed vs expected ratio, where uncertainty was assessed via bootstrapping at the genotype level where if a given genotype segment was sampled, the all other segments were also included to account for non-independence among segments within a genotype. We performed 1000 bootstrap replicates computing the mean observed/expected (O/E) and the 95% confidence interval. P-values were computed as the minimum proportion of replicates with O/E ratios above or below 1.

### The SIR with co-infection model

The susceptible-infected-recovered (SIR) compartmental model models the infection of the susceptible individuals by infected individuals, the recovery of infected individuals, and potentially waning immunity (SIRS model). We can simulate transmission histories under the SIR model, for example, by using the Gillespie algorithm gille-spie1977exact by keeping track of the parent-offspring relationships. The SIR model is parameterized by the transmission rate *θ*, the recovery rate *γ*, and the population size *N* . By default, the SIR model does not allow for individuals to be co-infected by multiple strains. However, we can easily account for this by modifying the transmission reaction such that infected individuals *I* can also infect *I* − 1 infected individuals (see also Figure S23.

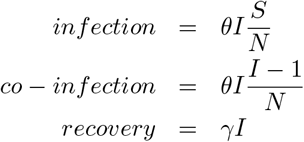

Conditional on a co-infection event, we assume that the probability of a reassortant virus within the co-infected individual is given by a constant c (see Figure S23). Upon recovery, there is a probability of that individual being sampled, given by the sampling proportion. The SIR with co-infection model follows the same S, I, and R dynamics as the SIR model without co-infection. By keeping track of the parent-offspring relationships, we can simulate transmission networks using the SIR with co-infection model.

We implemented two additional variations of the SIR with the co-infection model. First, we include super spreading by modeling the number of new infections given an infection event using a negative binomial distribution with mean 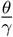 and variance 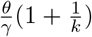, following the notation in, e.g. li2017quantifying. Second, simulate SIR with co-infection and superspreading dynamics across different sub-populations. To do so, we model *n* different populations with separate S, I, and R, between which infected individuals move at a rate given by the migration rate. We implemented simulators for SIR with co-infection, superspreading, and population structure into the BEAST2 package CoalRe.

### Connecting SIR models and the Coalescent with Reassortment

In the coalescent with reassortment, we model the co-evolution of multiple segments from the present to the past. Going back in time, lineages encounter reassortment events at a rate given by the reassortment rate *ρ*. Pairs of lineages can coalesce at a rate inversely proportional to the effective population size *N*_*e*_. The effective population size at time *t N*_*e*_(*t*), can be interpreted as a function of the parameters in the SIR model, as previously derived in volz2009phylodynamics and volz2012complex The effective population size can be denoted as:

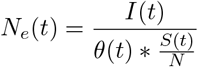

with *θ* (*t*) being the transmission rate at time t, *S*(*t*) being the number of susceptible individuals at time t, and *N* the total population size.

Similarly, we can express the reassortment rate *ρ* from the SIR with the co-infection model. In the forward in time SIR with co-infection model, the rate of co-infection between any two individuals at time *t* is given by 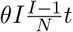. To convert this into a backward-in-time reassortment rate for the coalescent model, we can ask what the rate at which we expect to observe a reassortment event tracing a lineage backward in time from present to past is. First, we say the rate of reassortment on a specific lineage (the one we are tracking back in time) is:

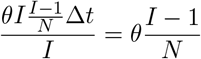

In the absence of any selection or preferential reassorting between specific lineages, we can assume that the creation of reassortant viruses is independent of time. We can therefore multiply the co-infection rate by the constant *c* to account for only a subset of co-infection events leading to reassortant viruses, to get the following expression for *ρ* (*t*)

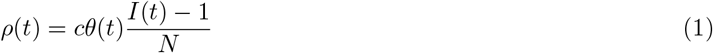

Unlike for the calculation of *N*_*e*_(*t*), there is no contribution of the susceptible fraction to the reassortment rates. This rate can be decomposed to model reassortment between distinct viral variants: when tracking the history of variant A, the rate of reassortment with variant B scales with the prevalence of B, since co-infection requires contact with B-infected hosts. Lastly, reassortant lineages are only detectable if at least one segment originates from a different parental lineage than all the other segments. As such, we can express the rate of observable reassortment events as:

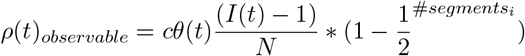

### The coalescent with reassortment closely approximates SIR with co-infection dynamics

To validate the above relationships between the SIR model and the Coalescent with Reassortment, we performed simulations under an SIR model with co-infection with 1000 individuals and a transmission rate of 0.1. We then simulated a coalescent with reassortment model with the same parameters and compared the results. We simulated 1000 replicates of each model, and compared the mean and variance of the Ne and *ρ* estimates. The wait time distribution between reassortment events is exponentially distributed with a mean of:

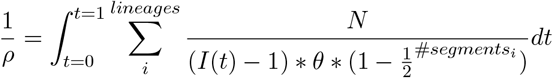

We find that the wait time distribution times the rate of reassortment events is exponentially distributed with mean 1 (see Figure S27A). When we add superspreading to the simulations using a Poisson-distributed number of infection events, the true wait time distribution includes 0, as there can be multiple co-infection events occurring at the same time, which is not modeled by our formulation of the reassortment rate (see Figure S27B). Importantly, this formulation of the reassortment rates is independent of any sampling process.

Lastly, we show how the reassortment rate across different locations can be defined as a function of the weighted average across these locations. To do so, we state that

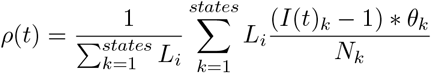

Which describes the reassortment rates at time *t* as a weighted average of the reassortment rates across all sub-populations *k*, weighted by the number of lineages in states *k* at time *t*. In other words, we can directly express the reassortment rates across an arbitrary number of states as the weighted average of the reassortment rates of each state. Importantly, this is not the case for other quantities sometimes used to approximate disease prevalence, such as the effective population size Ne. We show in Figure S27C that, given knowledge of lineages, the prevalence of the different states and transmission rates allows us to directly compute the reassortment rate.

### Implementation

To accurately model reassortment dynamics, we implemented time-varying reassortment rates in the BEAST2 package CoalRe. We first define a time grid, which is provided as a model input. At each grid point, we estimate the log reassortment rate, log *ρ* (*t*). Between grid points, we assume that log *ρ* (*t*) changes linearly through time, which corresponds to assuming exponential growth or decline of *ρ* (*t*) between grid points. This approach is analogous to how effective population size trajectories are modeled in phylodynamic methods such as the Skygrid model [17].

As a prior on log *ρ* (*t*), we use a Gaussian Markov random field (GMRF), such that

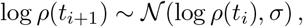

which encourages temporal smoothness while allowing flexible changes in reassortment rates through time. We model log *N*_*e*_(*t*) analogously using the same GMRF framework. We further allow the reassortment rate to depend on other time-varying quantities:

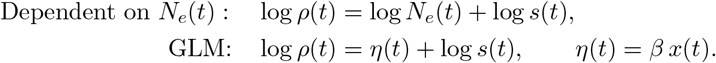

Here, *ρ* (*t*) denotes the reassortment rate at time *t*, and *N*_*e*_(*t*) is the effective population size. The term *s*(*t*) is a positive scaling function that captures deviations in reassortment intensity not explained by *N*_*e*_(*t*) or other predictors and is treated as an offset on the log scale.

In the GLM formulation, *x*(*t*) represents a time-varying predictor (e.g., a covariate or spline-based function of time), and *β* is a regression coefficient quantifying the strength and direction of its association with reassortment rates. The linear predictor *η* (*t*) allows reassortment dynamics to be flexibly linked to external covariates while retaining the GMRF prior on temporal smoothness.

### Validation of the Coalescent with time-varying reassortment

To validate the implementation, we first compared the networks simulated under the coalescent with a time-varying coalescent prior that describes the same distributions as networks sampled under the same prior using Markov chain Monte Carlo. This is an approach frequently used to validate model implementations. As shown in Figure S28, networks sampled and simulated under the same model describe the same distribution.

We next tested the implementation by simulating networks under SIR with infection models, SIR with co-infection and superspreading, and structured SIR with superspreading models. For each model, we performed 10 simulations. We then simulate genomic sequence data using seq-gen for each segment. For each simulation, we then performed three different inferences. First, we assumed constant reassortment rates (constant; Figure S29). Next, we modelled the *ρ* (*t*) as a time-varying function (skygrowth; Figure S30). Lastly, we modelled *ρ* (*t*) as a function of *N*_*e*_(*t*) (skygrowth Ne).

We show in Figure S28, Figure S31, Figure S34, and Figure S35 that we can accurately reconstruct disease prevalences from reassortment rates when the transmission rates are known. The validation results demonstrate close agreement between simulated and inferred reassortment dynamics across all model scenarios (Figure S36, Figure S32, Figure S33). These results confirm that our method can infer epidemiological parameters from reassortment networks.

## Code Availability

All code for the described analyses and visualization of results can be found in the following GitHub repository: https://github.com/nicfel/ReassortmentDynamics. Sequences were simulated using Seq-Gen [34], phylogenetic trees and networks were visualized using an adapted version of baltic https://github.com/evogytis/baltic, and plots were generated using matplotlib [20] and ggplot2 [41].

## Data Availability

Sequence data were obtained from the GISAID EpiFlu database (https://gisaid.org)m, accession identifiers and acknowledgments of submitting laboratories are provided in the Supplementary Materials.

## Acknowledgments

This work was funded by the Center for Forecasting and Outbreak Analytics of the US Centers for Disease Control & Prevention via the Insight Net cooperative agreement (CDC-RFA-FT-23-0069 to SYT and JAL). Its contents are solely the responsibility of the authors and do not necessarily represent the official views of the Centers for Disease Control and Prevention. We would like to thank Adrian Castellanos and Barbara Han for help sourcing LPAI case data from the APHIS dashboard.

