## Supplemental File for "Frequent seasonal reassortment between high and low path viruses drives the diversification of influenza A/H5N1"

### Supplementary material

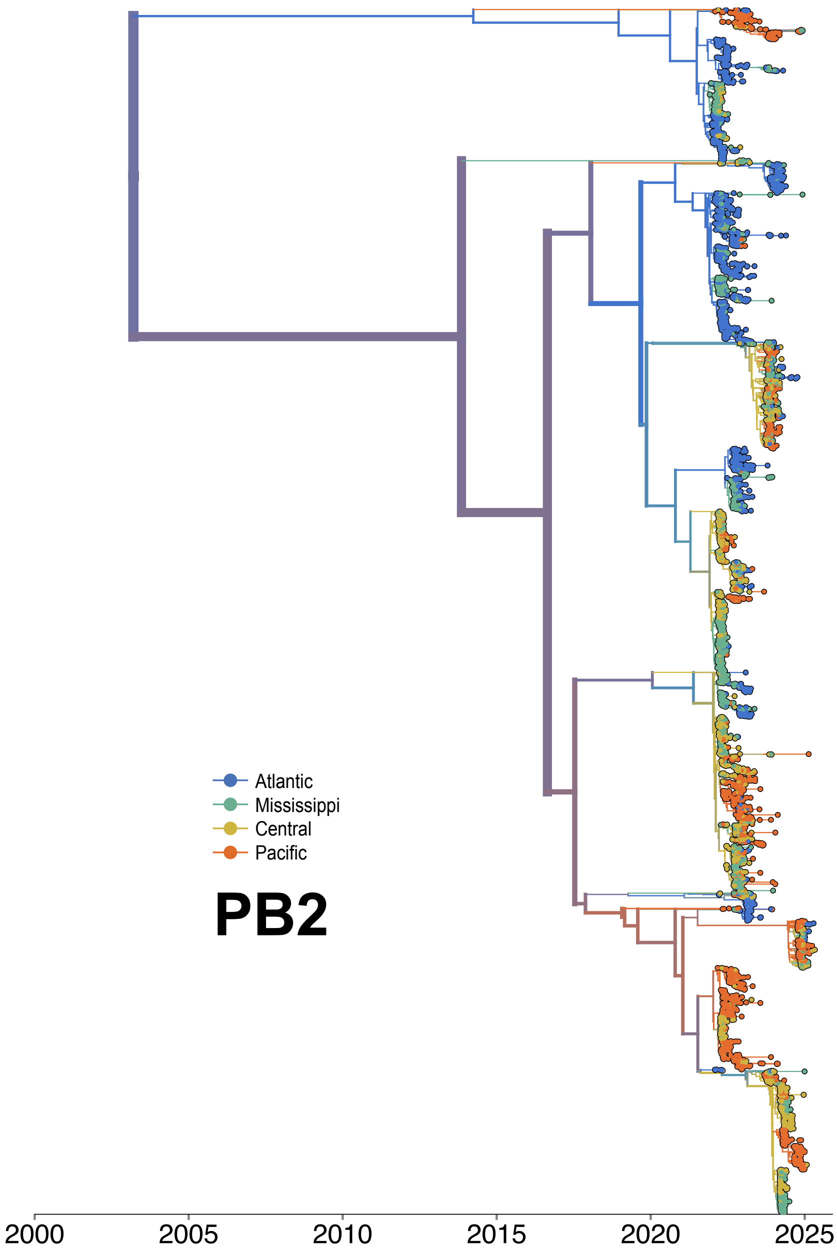


Figure S1: **MCC tree for PB2 segment colored by migratory flyway.** Maximum Clade Credibility tree for 9052 HPAI H5N1 sequences from North America with branches colored by migratory flyway inferred using discrete trait diffusion modeling.

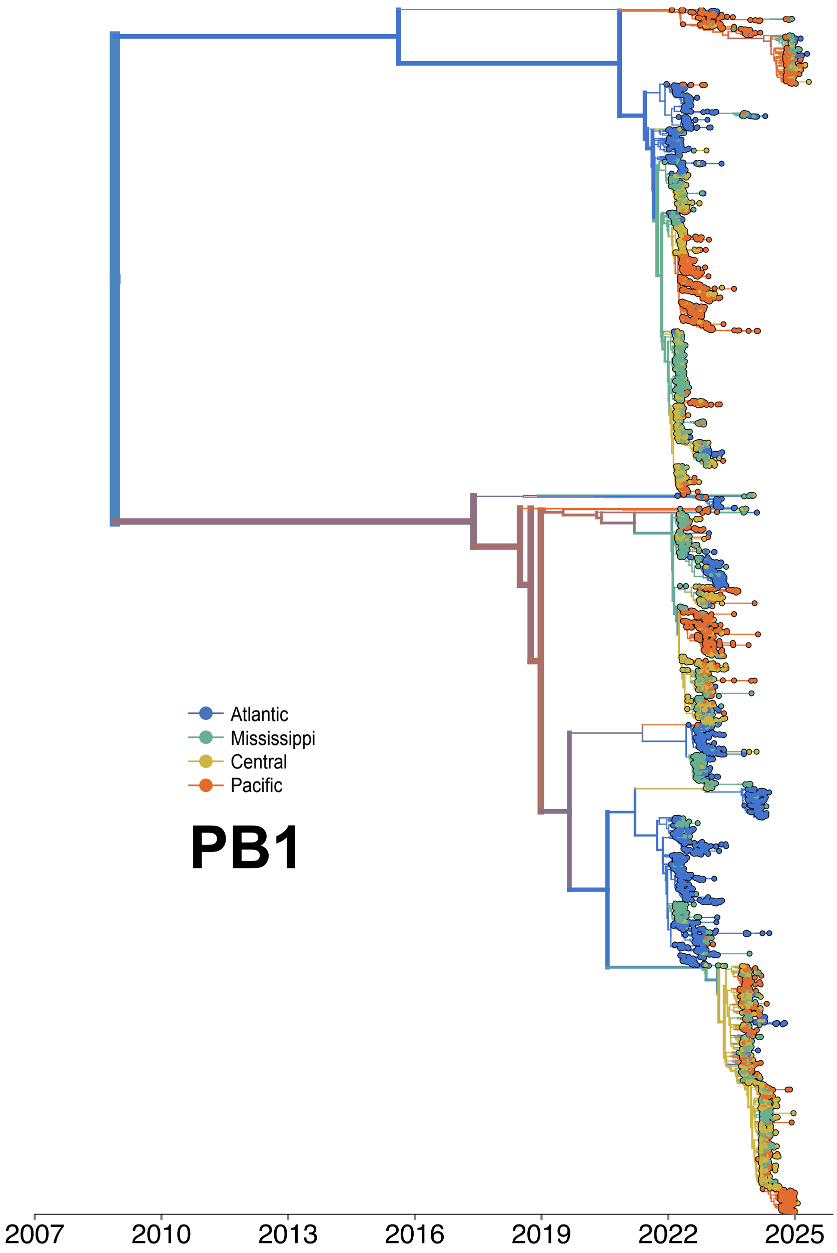


Figure S2: **MCC tree for PB1 segment colored by migratory flyway.** Maximum Clade Credibility tree for 9052 HPAI H5N1 sequences from North America with branches colored by migratory flyway inferred using discrete trait diffusion modeling.

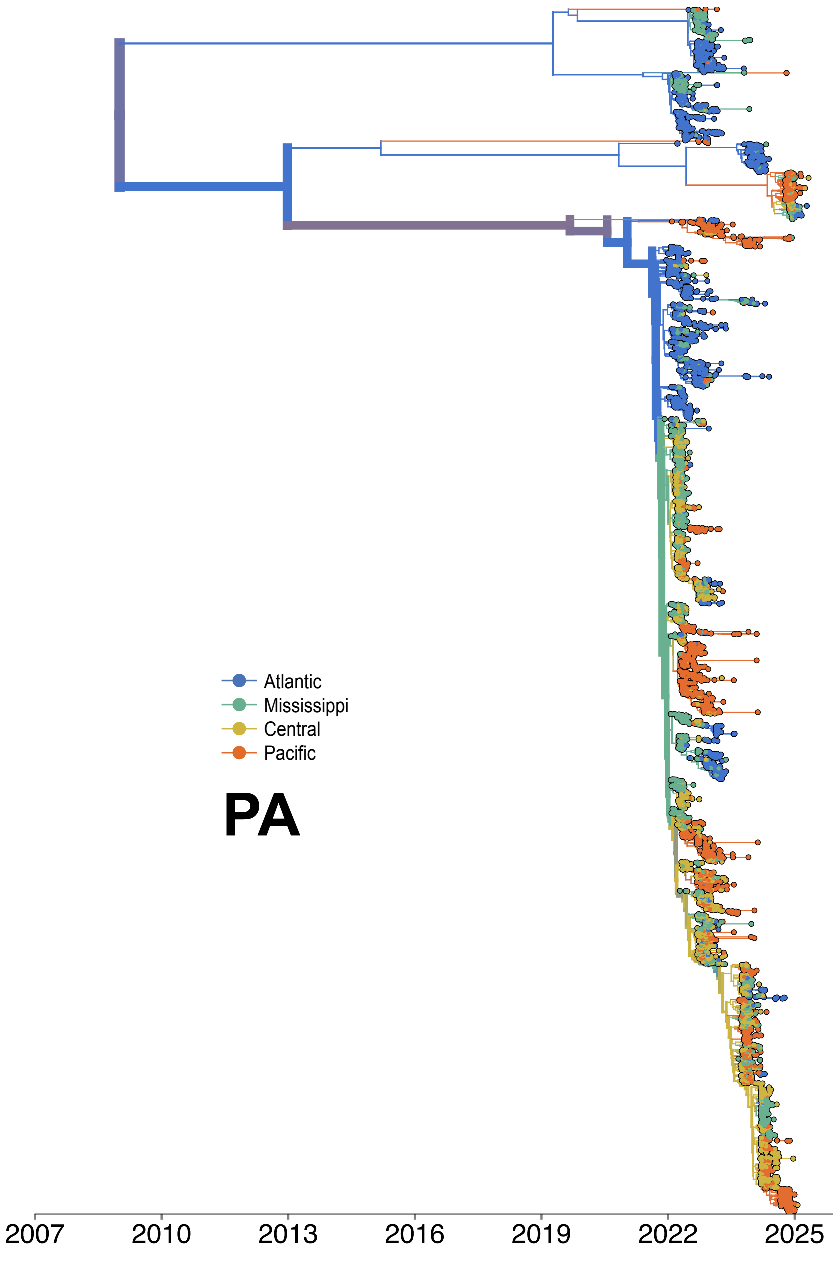


Figure S3: **MCC tree for PA segment colored by migratory flyway.** Maximum Clade Credibility tree for 9052 HPAI H5N1 sequences from North America with branches colored by migratory flyway inferred using discrete trait diffusion modeling.

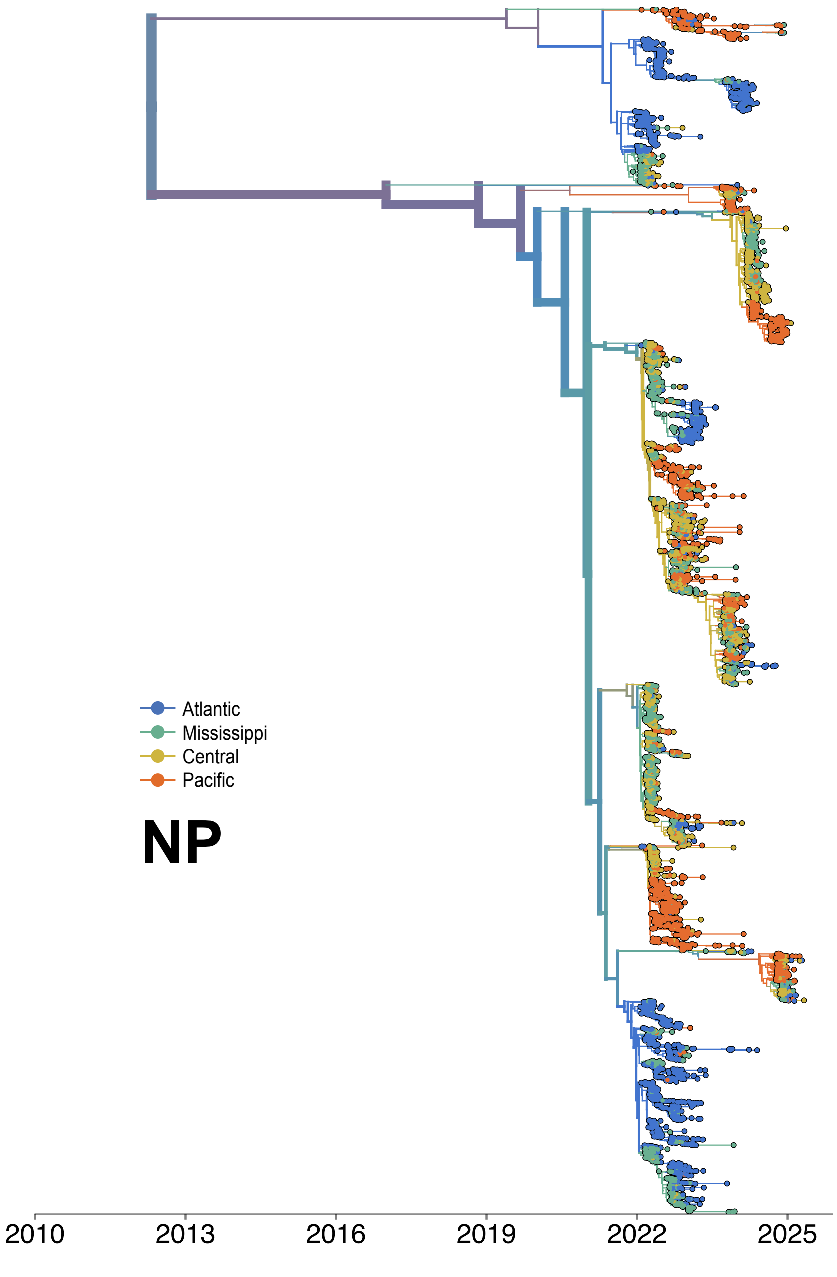


Figure S4: **MCC tree for NP segment colored by migratory flyway.** Maximum Clade Credibility tree for 9052 HPAI H5N1 sequences from North America with branches colored by migratory flyway inferred using discrete trait diffusion modeling.

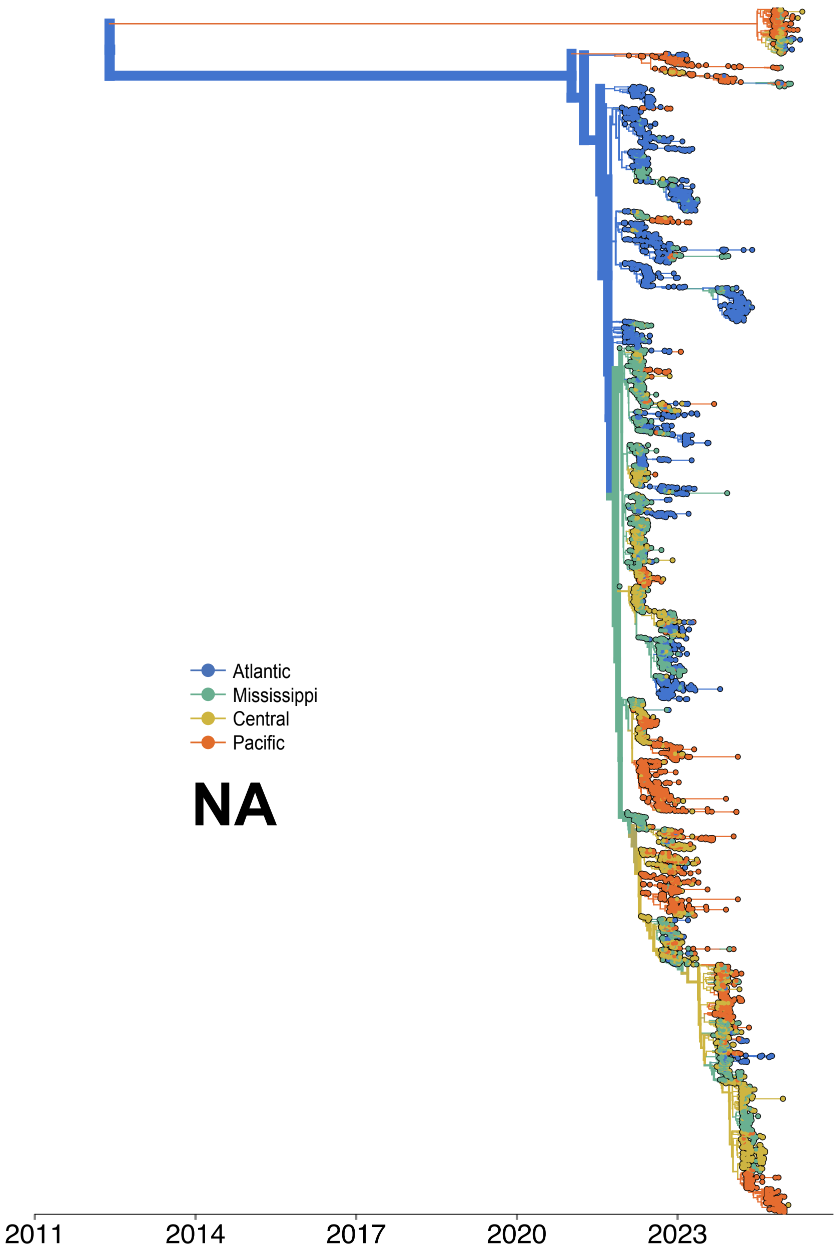


Figure S5: **MCC tree for NA segment colored by migratory flyway.** Maximum Clade Credibility tree for 9052 HPAI H5N1 sequences from North America with branches colored by migratory flyway inferred using discrete trait diffusion modeling.

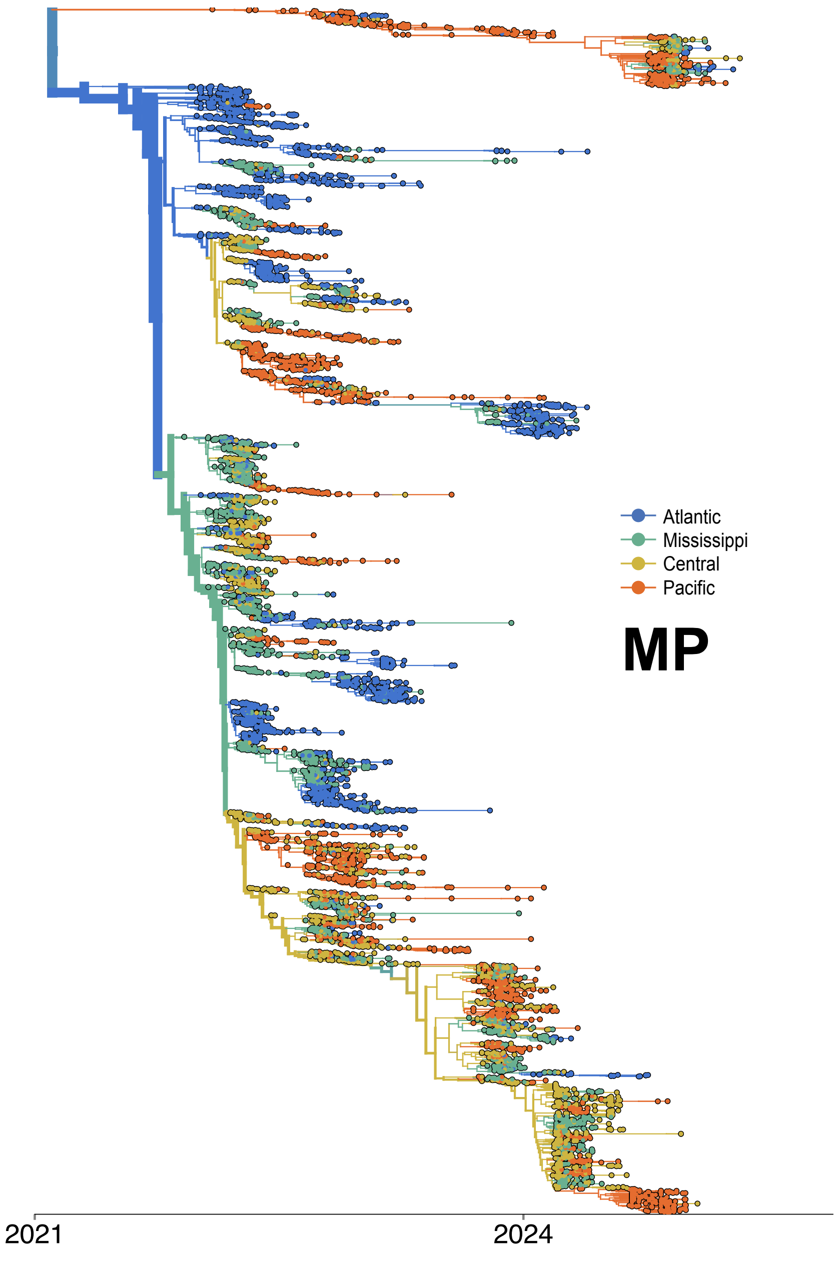


Figure S6: **MCC tree for MP segment colored by migratory flyway.** Maximum Clade Credibility tree for 9052 HPAI H5N1 sequences from North America with branches colored by migratory flyway inferred using discrete trait diffusion modeling.

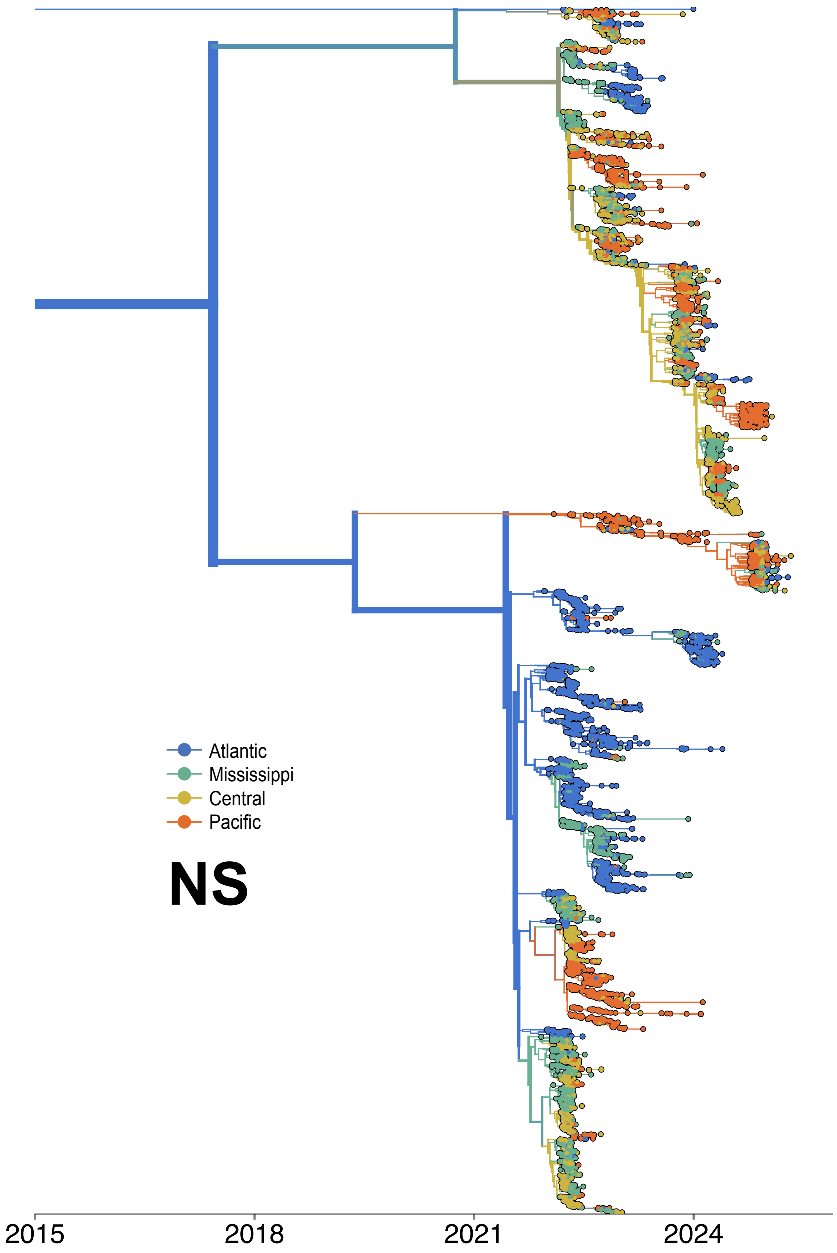


Figure S7: **MCC tree for NS segment colored by migratory flyway.** Maximum Clade Credibility tree for 9052 HPAI H5N1 sequences from North America with branches colored by migratory flyway inferred using discrete trait diffusion modeling.

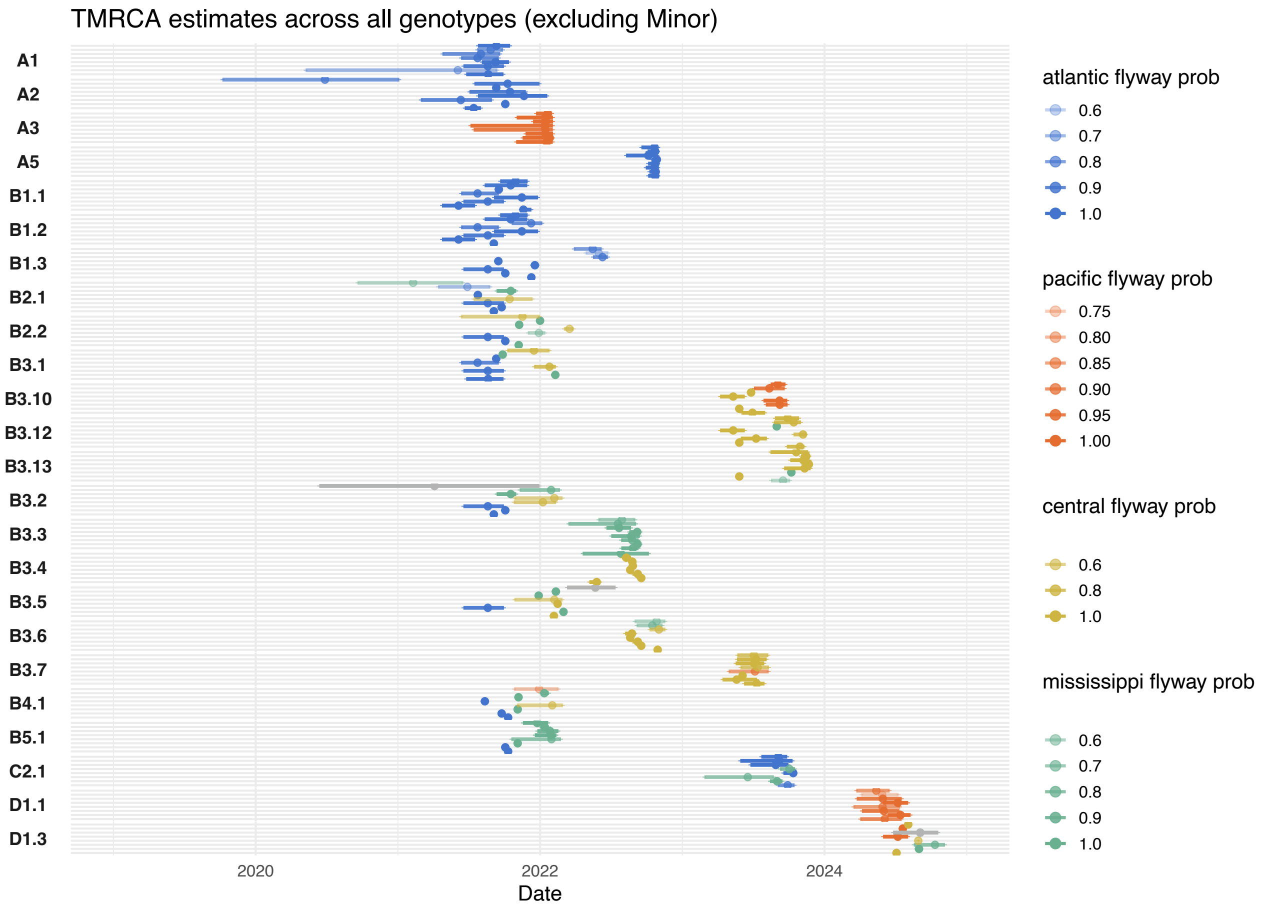


Figure S8: **TMRCA estimates by migratory flyway.** Time to most recent common ancestor estimates for each genoFLU genotype for each segment colored by inferred migratory flyway. The mean value is plotted with 95% HPD interval. Opacity of color corresponds to the posterior probability of the inferred migratory flyway; segments with posterior probability greater than 50% are colored grey.

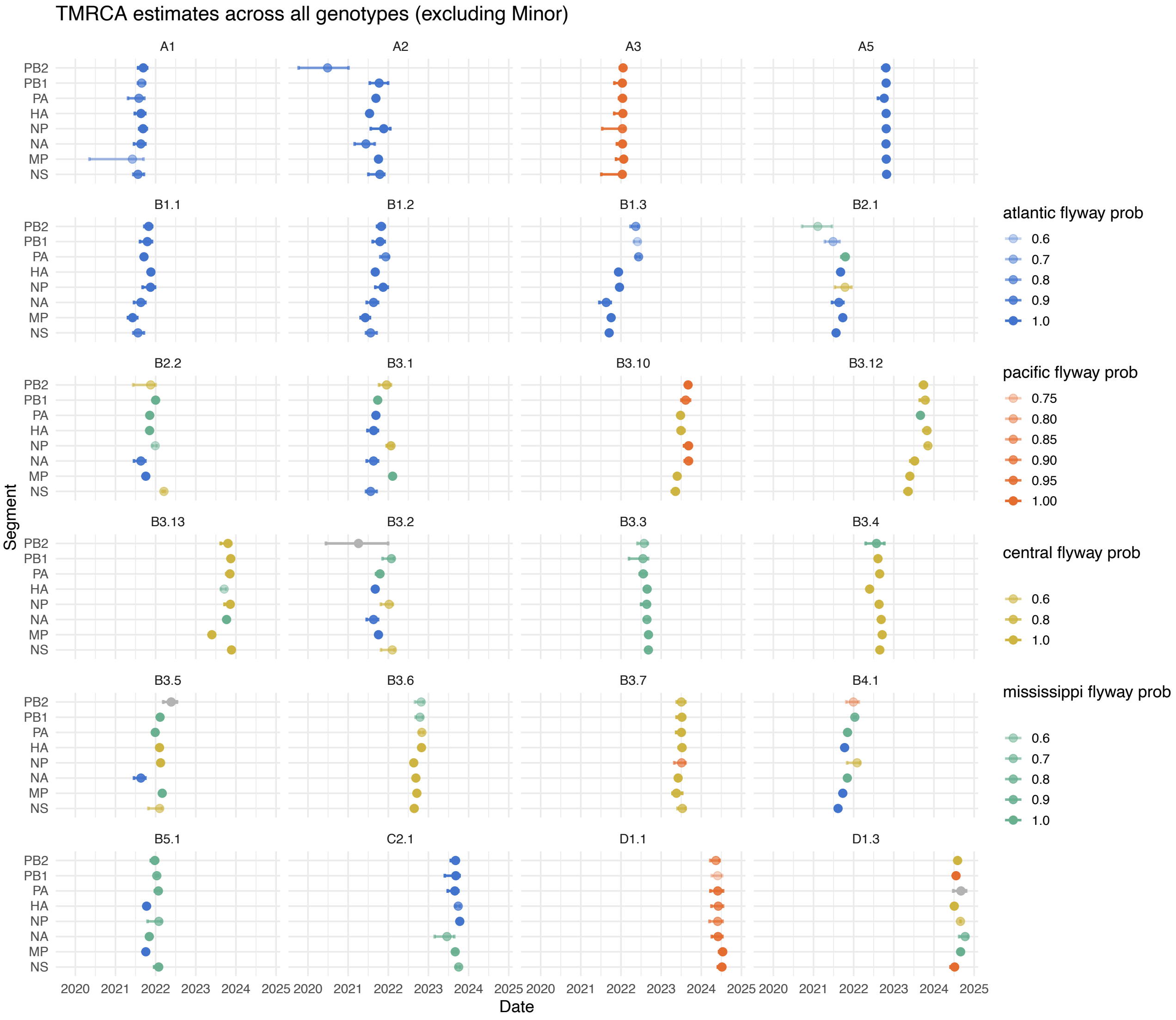


Figure S9: **TMRCA estimates by migratory flyway, faceted by genotype.** Time to most recent common ancestor estimates for each genoFLU genotype for each segment colored by inferred migratory flyway faceted by each genoFLU genotype. The mean value is plotted with 95% HPD interval. Opacity of color corresponds to the posterior probability of the inferred migratory flyway; segments with posterior probability greater than 50% are colored grey.

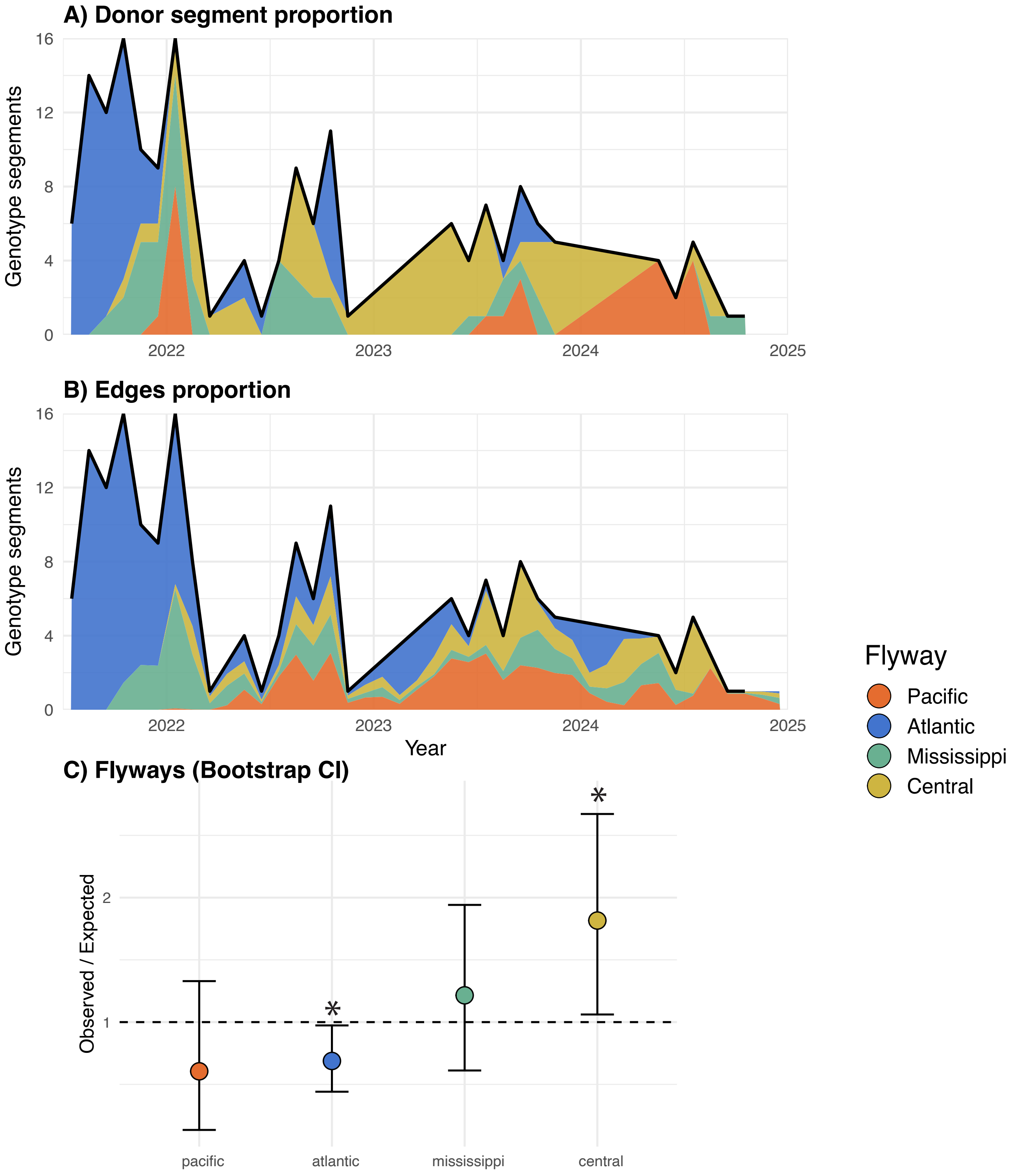


Figure S10: **Genotype emergence and transmission by host and flyway. A** Number of genotype segments in monthly time-bins with the proportion of segments inferred from a given host colored below the line. **B** Number of genotype segments in monthly time-bins with the proportion of edges across the phylogeny coming from a given host colored below the line. **C** Results of Observed/Expected ratio for genotype emergence from a given migratory flyway.

| Flyway | observed n | expected mean | oe_ratio_mean | oe_lower_boot | oe_upper_boot | p_boot | p_label | signif |
|---|---|---|---|---|---|---|---|---|
| Atlantic | 56 | 81.3364 | 0.687671536 | 0.440807881 | 0.973373848 | 0.031968032 | 3.20E-02 | * |
| Central | 48 | 26.9389 | 1.815768318 | 1.061207609 | 2.67172232 | 0.037962038 | 3.80E-02 | * |
| Mississippi | 38 | 31.4462 | 1.215009978 | 0.61140414 | 1.941367978 | 0.533466533 | 5.33E-01 | |
| Pacific | 22 | 38.2785 | 0.604757759 | 0.133784365 | 1.329203777 | 0.222 | 2.22E-01 | |

Figure S11: **Results of permutation test for migratory flyways.**

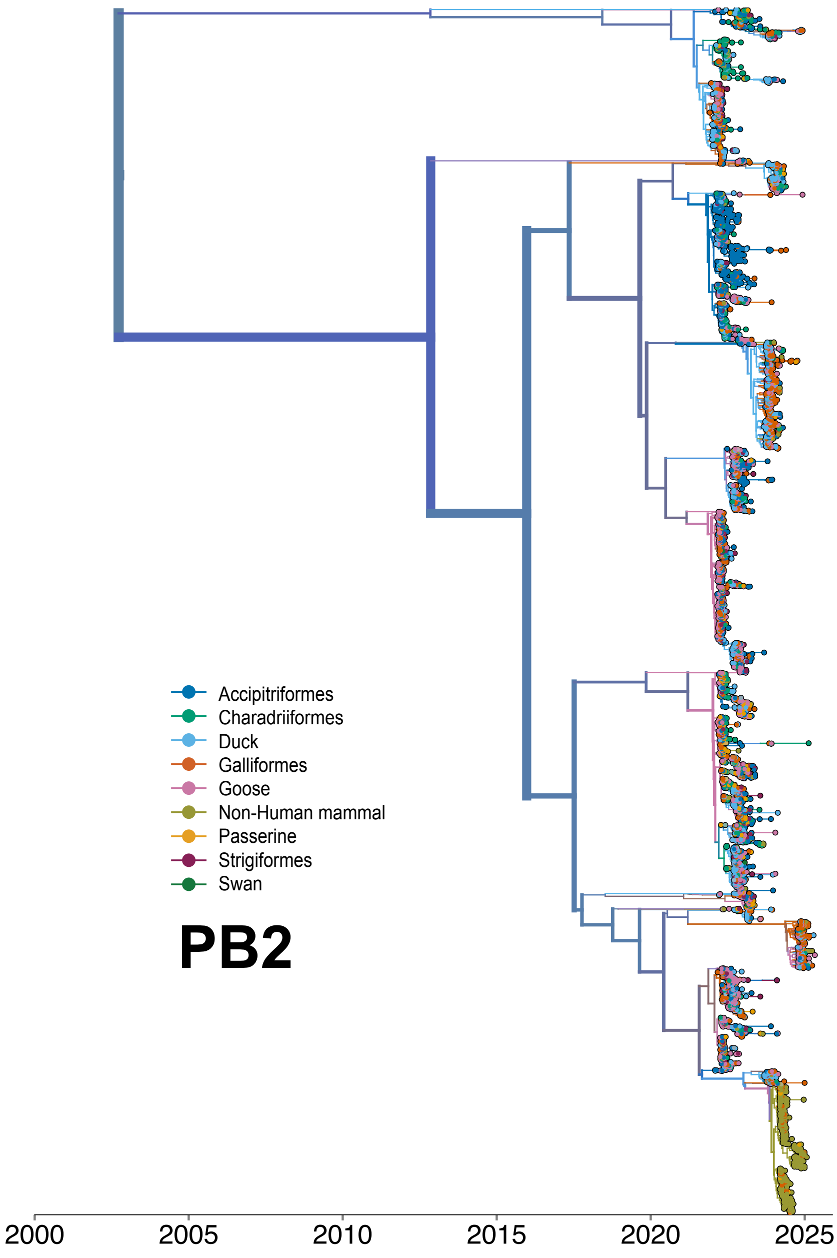


Figure S12: **MCC tree for PB2 segment colored by host.** Maximum Clade Credibility tree for 9052 HPAI H5N1 sequences from North America with branches colored by host inferred using discrete trait diffusion modeling.

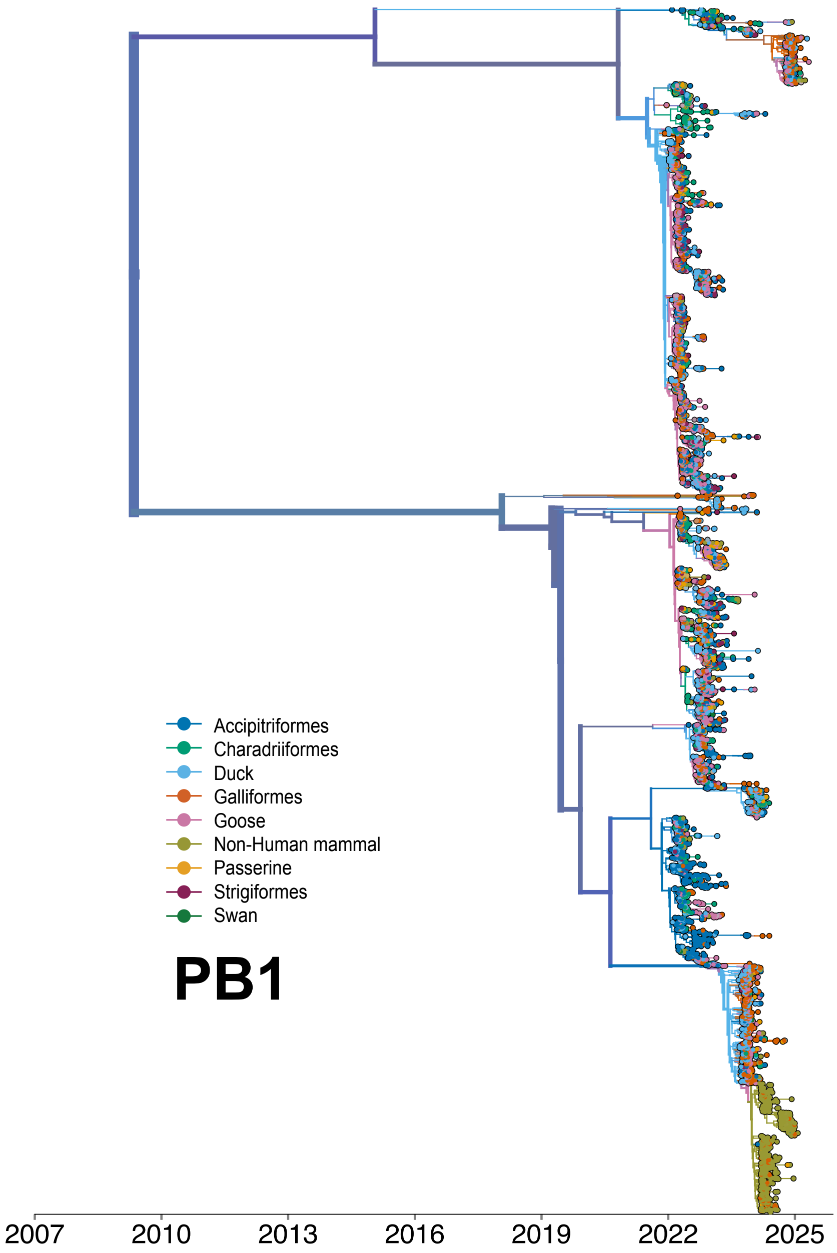


Figure S13: **MCC tree for PB1 segment colored by host.** Maximum Clade Credibility tree for 9052 HPAI H5N1 sequences from North America with branches colored by host inferred using discrete trait diffusion modeling.

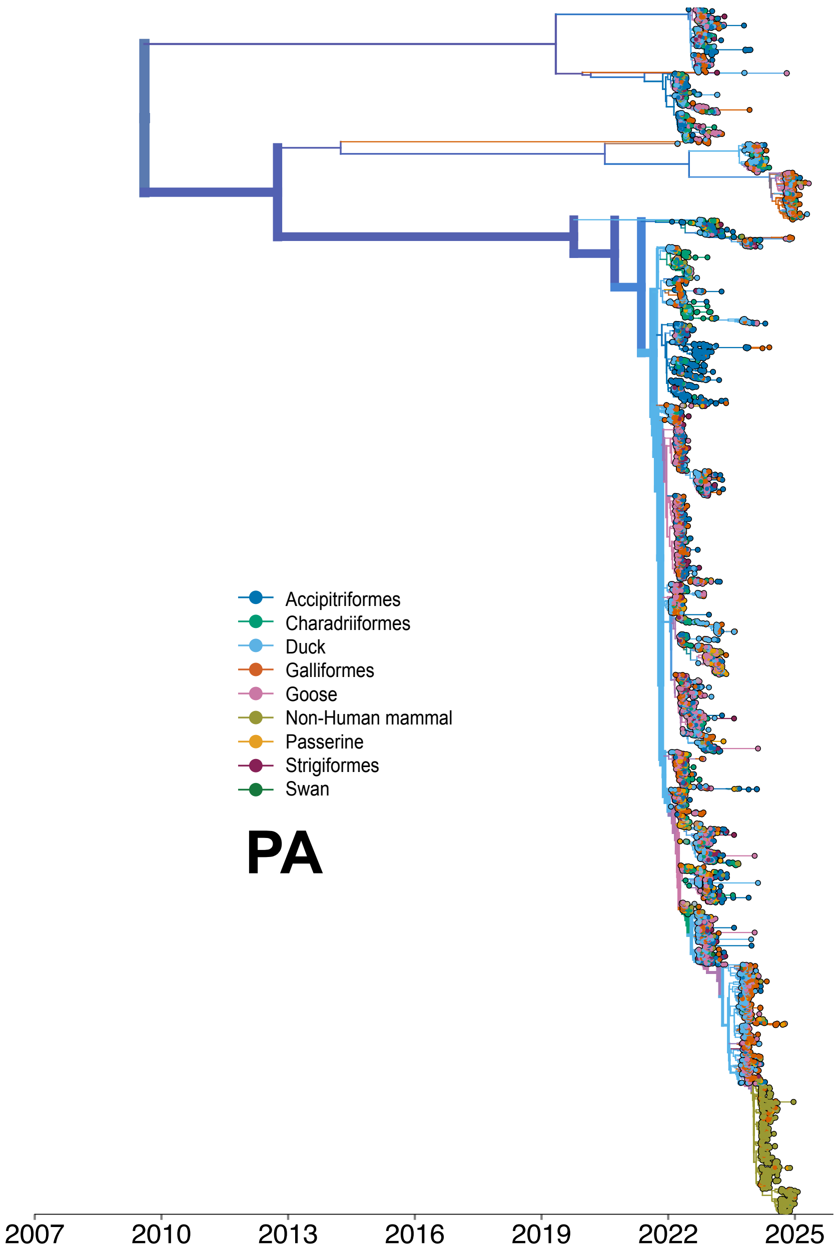


Figure S14: **MCC tree for PA segment colored by host.** Maximum Clade Credibility tree for 9052 HPAI H5N1 sequences from North America with branches colored by host inferred using discrete trait diffusion modeling.

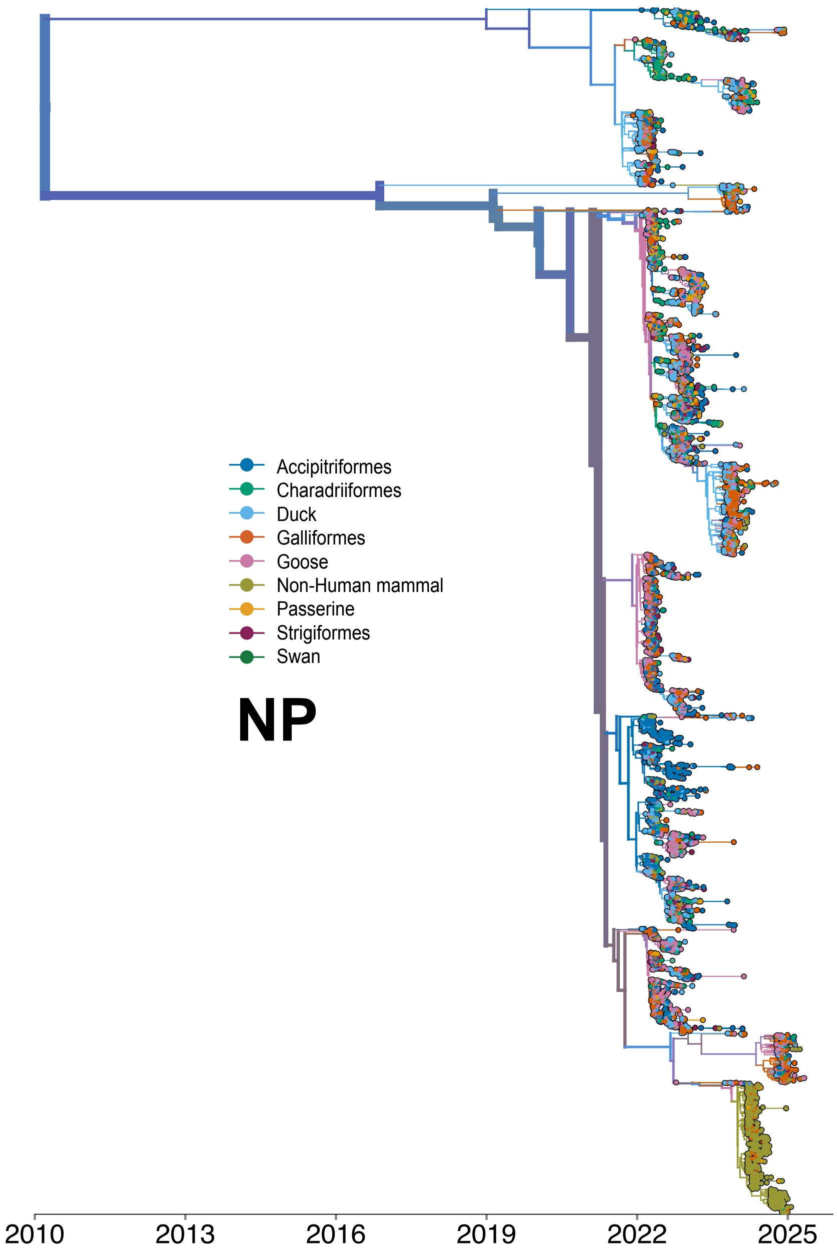


Figure S15: **MCC tree for NP segment colored by host.** Maximum Clade Credibility tree for 9052 HPAI H5N1 sequences from North America with branches colored by host inferred using discrete trait diffusion modeling.

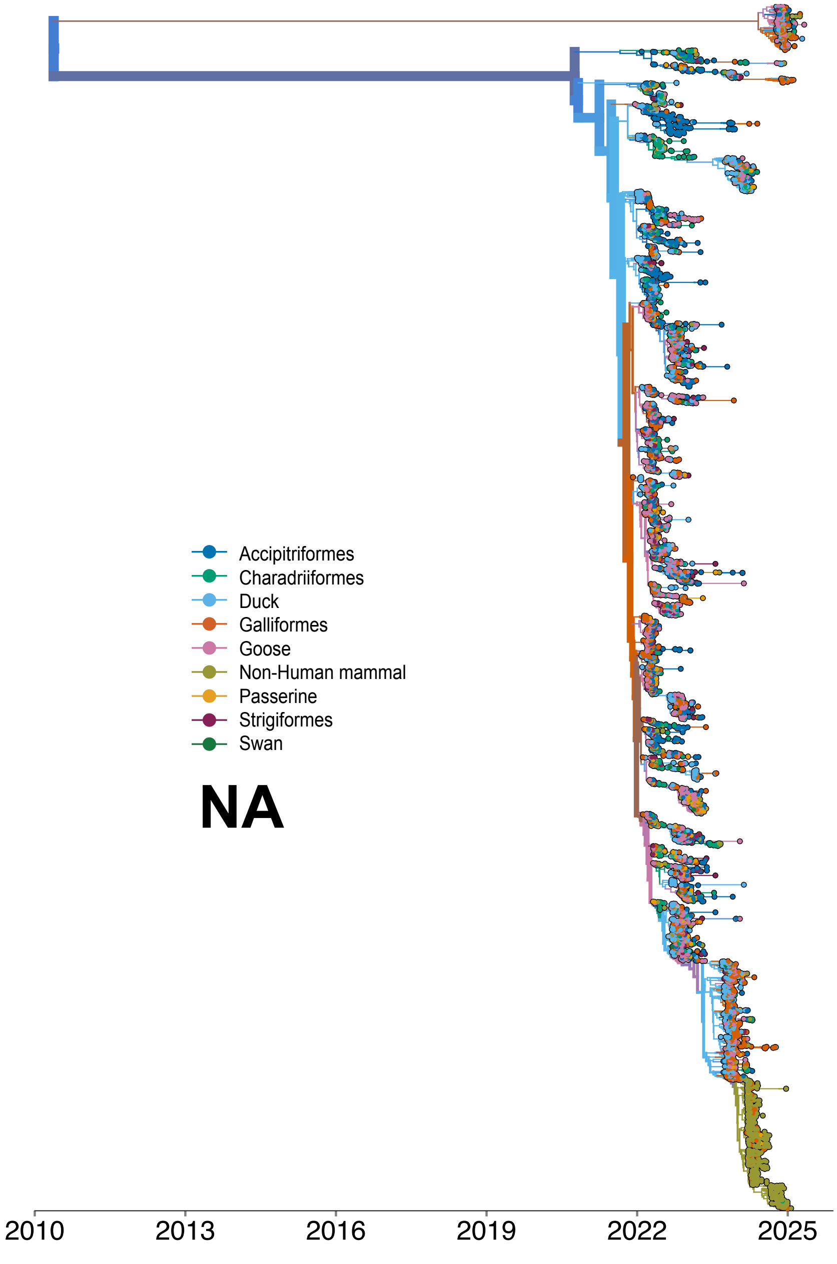


Figure S16: **MCC tree for NA segment colored by host.** Maximum Clade Credibility tree for 9052 HPAI H5N1 sequences from North America with branches colored by host inferred using discrete trait diffusion modeling.

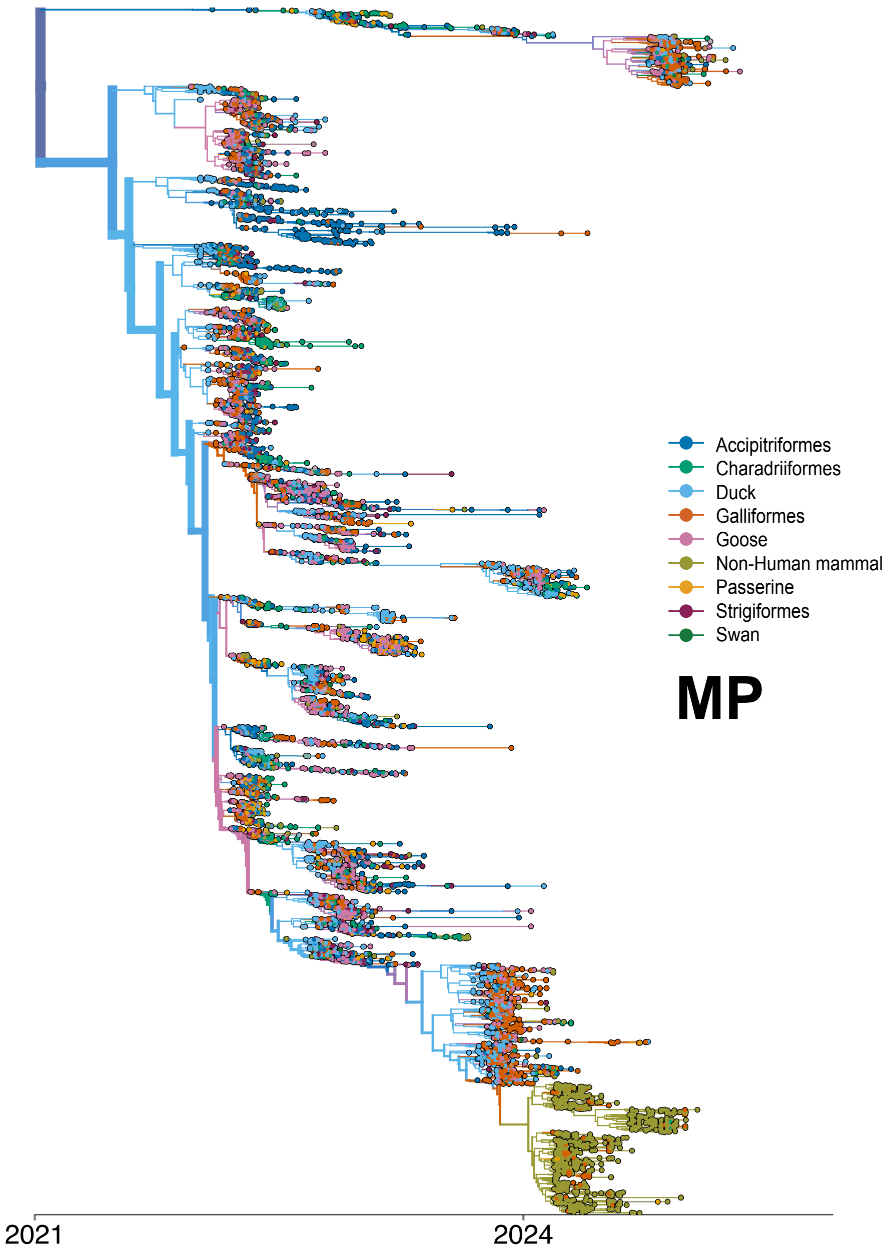


Figure S17: **MCC tree for MP segment colored by host.** Maximum Clade Credibility tree for 9052 HPAI H5N1 sequences from North America with branches colored by host inferred using discrete trait diffusion modeling.

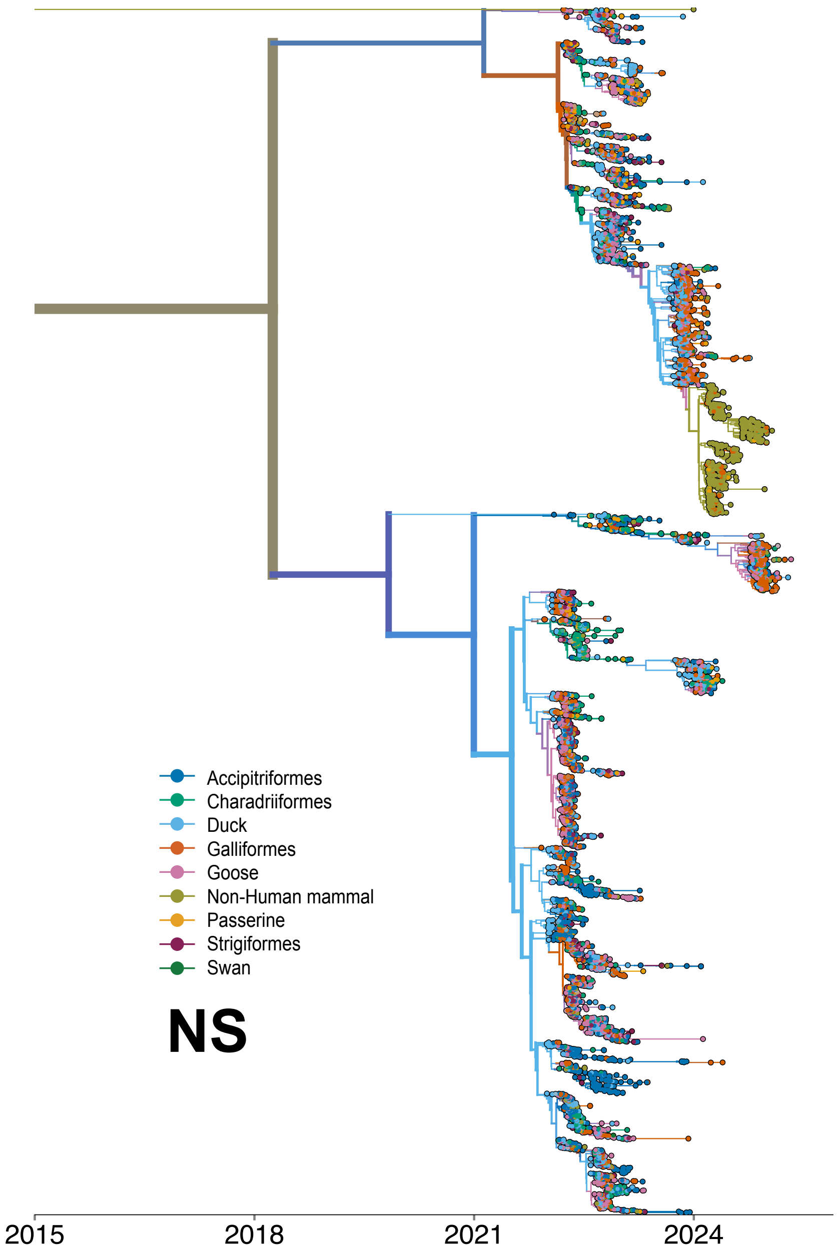


Figure S18: **MCC tree for NS segment colored by host.** Maximum Clade Credibility tree for 9052 HPAI H5N1 sequences from North America with branches colored by host inferred using discrete trait diffusion modeling.

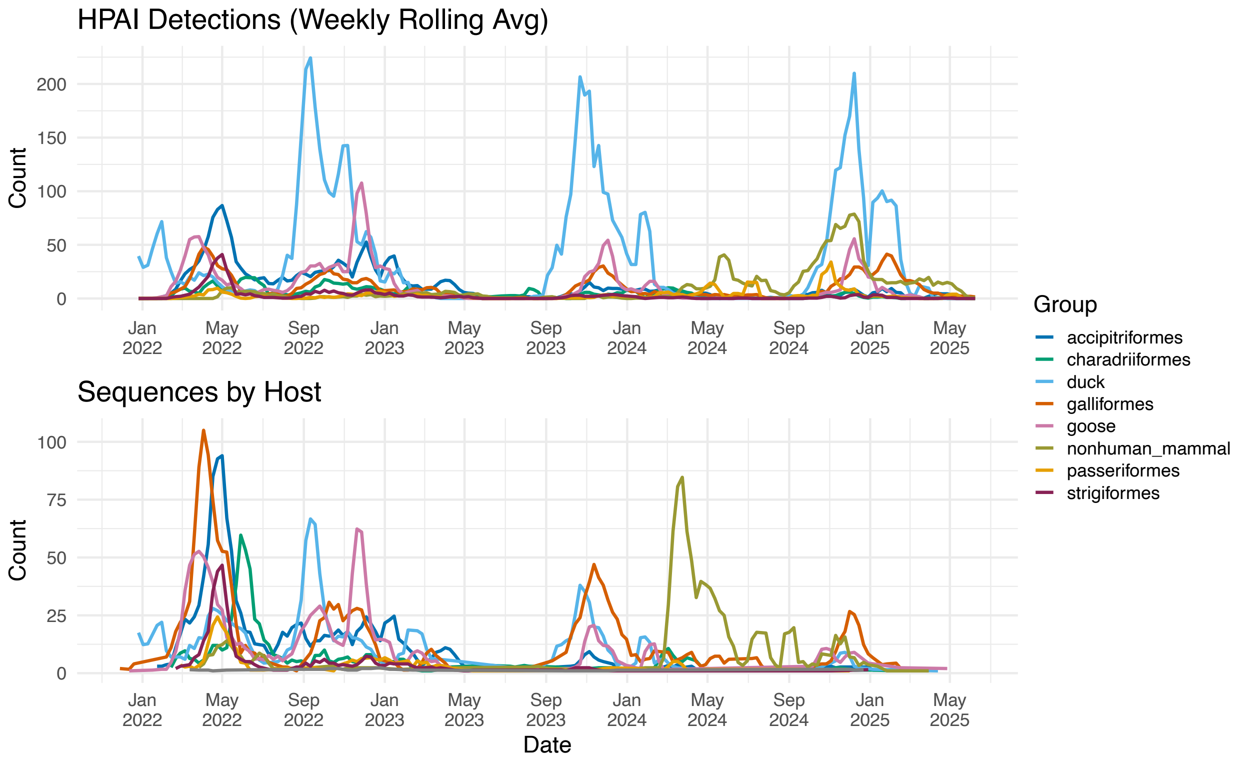


Figure S19: **Detections and sequences by host group.** Number of detections and sequences for HPAI H5N1 in North America by host group 2022-2025.

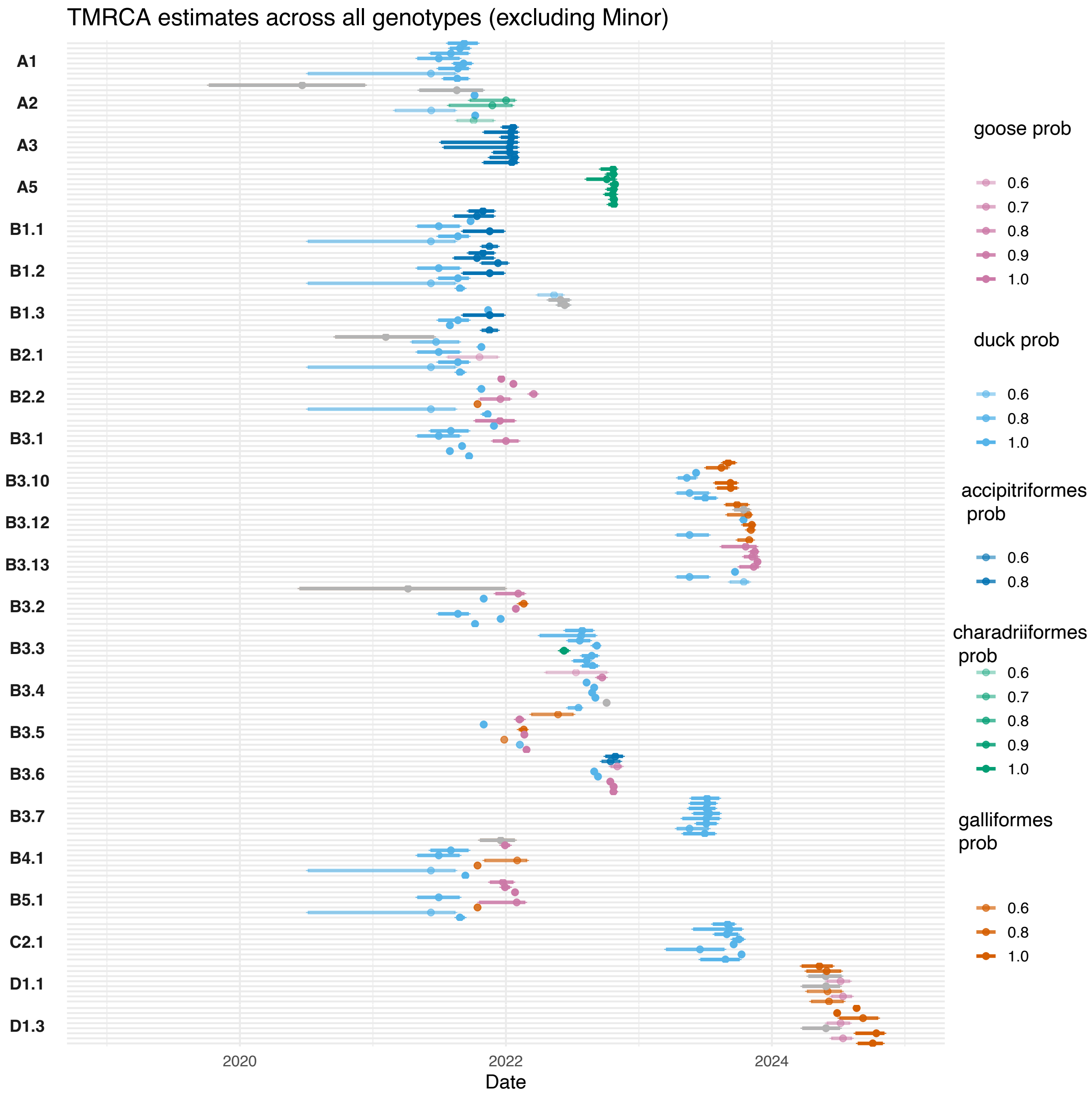


Figure S20: **TMRCA estimates by host.** Time to most recent common ancestor estimates for each genoFLU genotype for each segment colored by inferred host. The mean value is plotted with 95% HPD interval. Opacity of color corresponds to the posterior probability of the inferred host; segments with posterior probability greater than 50% are colored grey.

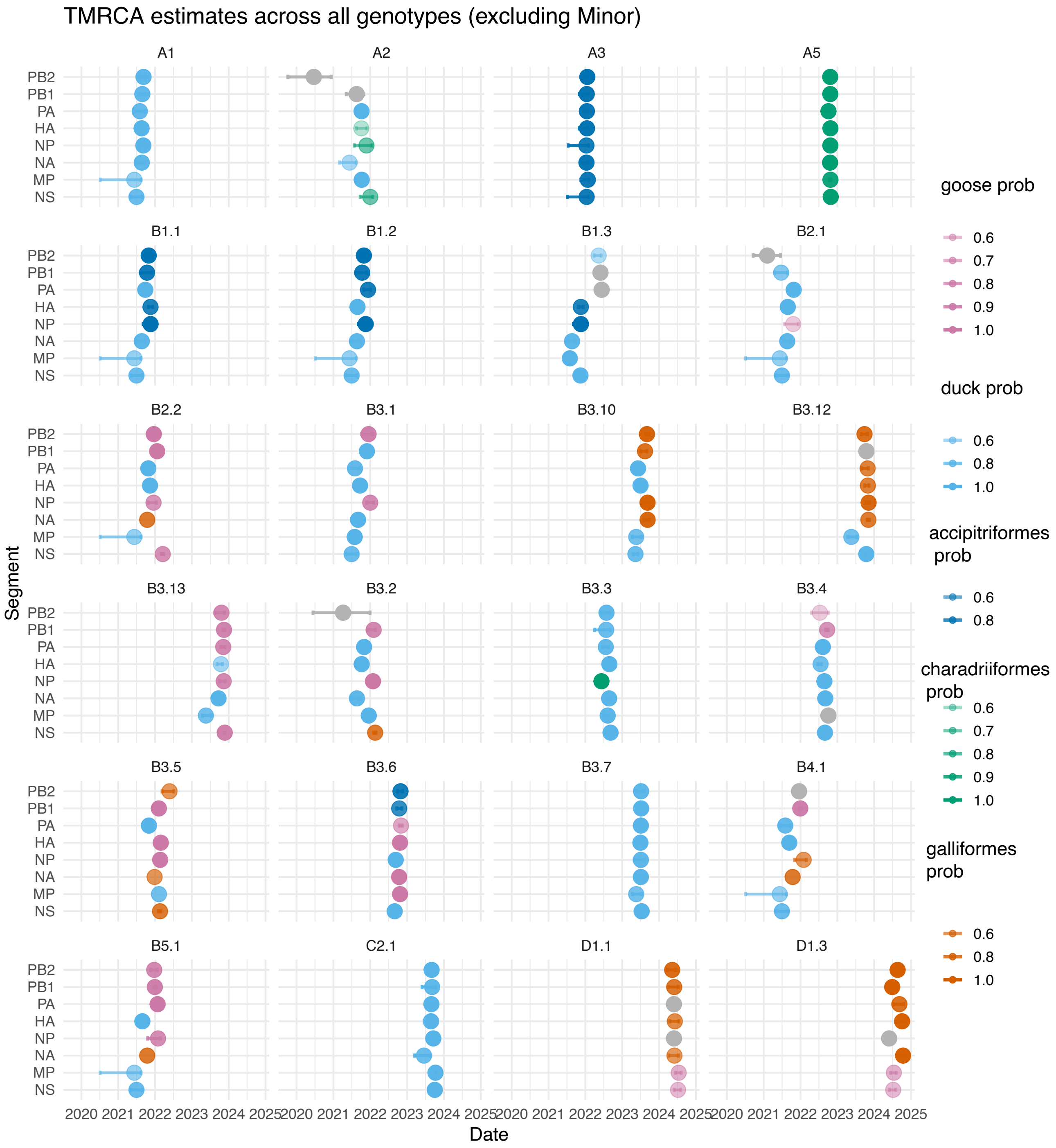


Figure S21: **TMRCA estimates by host, faceted by genotype.** Time to most recent common ancestor estimates for each genoFLU genotype for each segment colored by inferred host faceted by each genoFLU genotype. The mean value is plotted with 95% HPD interval. Opacity of color corresponds to the posterior probability of the inferred host; segments with posterior probability greater than 50% are colored grey.

| group | observed n | expected mean | oe_ratio_mean | oe_lower_boot | oe_upper_boot | p_boot | p_label | signif |
|---|---|---|---|---|---|---|---|---|
| accipitriformes | 17 | 16.1472 | 1.059655478 | 0.235228468 | 2.150459304 | 0.959839357 | 9.60E-01 | |
| charadriiformes | 11 | 10.5797 | 1.134212432 | 0.113556261 | 3.396820413 | 0.966597077 | 9.67E-01 | |
| duck | 80 | 98.0954 | 0.818481311 | 0.619182425 | 1.036916334 | 0.125874126 | 1.26E-01 | |
| galliformes | 24 | 20.6551 | 1.197884514 | 0.481327792 | 1.952056066 | 0.599400599 | 5.99E-01 | |
| goose | 32 | 16.0633 | 2.028170458 | 1.038139371 | 3.319881952 | 0.041958042 | 4.20E-02 | * |

Figure S22: **Results of permutation test for host.**

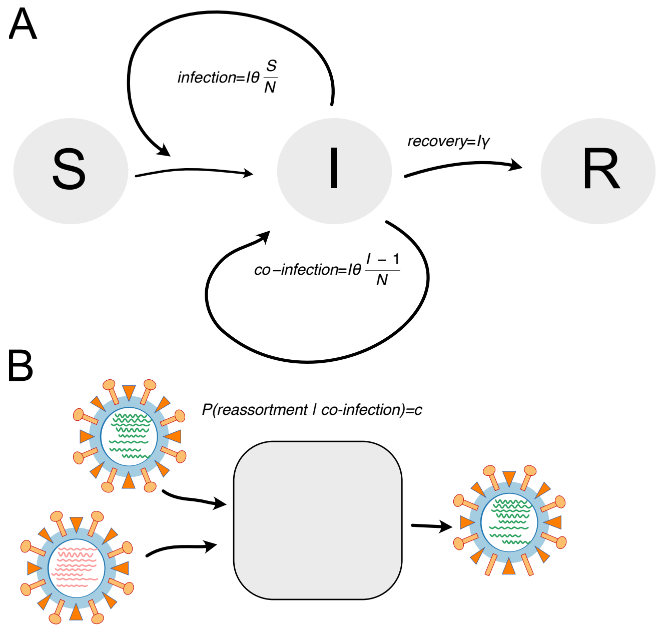


Figure S23: **Principle of the SIR with co-infection model. A** The SIR with the co-infection model allows modeling transmission dynamics using classical SIR models while adding a co-infection reaction. The rate of co-infection is given by the transmission rates times the number of infected individuals in a population. For each infection event, there is an $\frac{I-1}{N}$ probability that the infection event is with another infected individual. **B** Conditional on an event where an infected individual is subject to another infection event, reassortment of the two viral lineages can occur. The probability of reassortment conditional on co-infection is dependent on, for example, superinfection exclusion and the time difference between the two infection events. We here simplify this probability into a constant c.

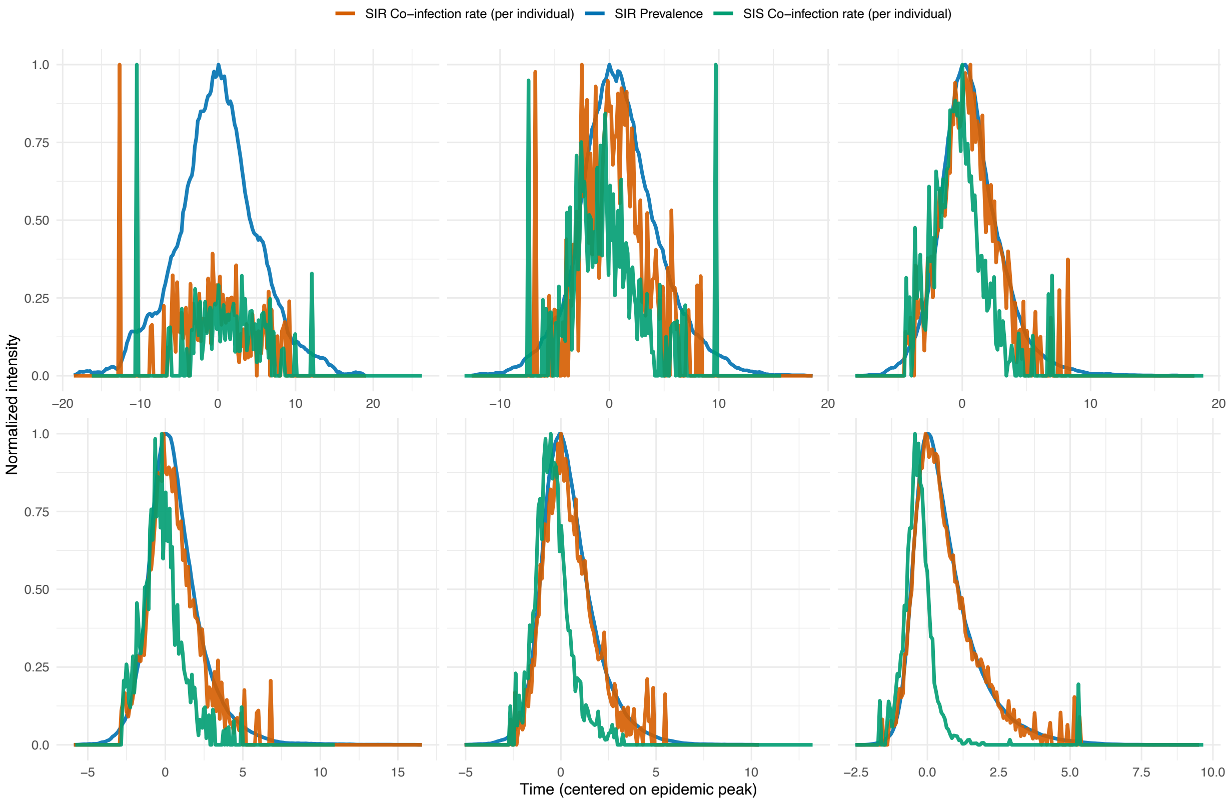


Figure S24: **Comparison of prevalence and co-infection rate dynamics between SIR and SIS models.** Time series comparison of normalized prevalence and co-infection rates across different reproduction numbers ($R_0$). Blue lines show SIR prevalence dynamics, red lines show SIR co-infection rates, and green lines show SIS co-infection rates. All metrics are normalized to their maximum value within each $R_0$ scenario to enable comparison of temporal patterns. Time is centered on the epidemic peak to align dynamics across different $R_0$ values. Panels show results for $R_0$ ranging from low to high transmission scenarios. The comparison reveals how co-infection opportunities vary between SIR (susceptible-infected-recovered) and SIS (susceptible-infected-susceptible) transmission models, with important implications for reassortment dynamics in populations with different immunity patterns.

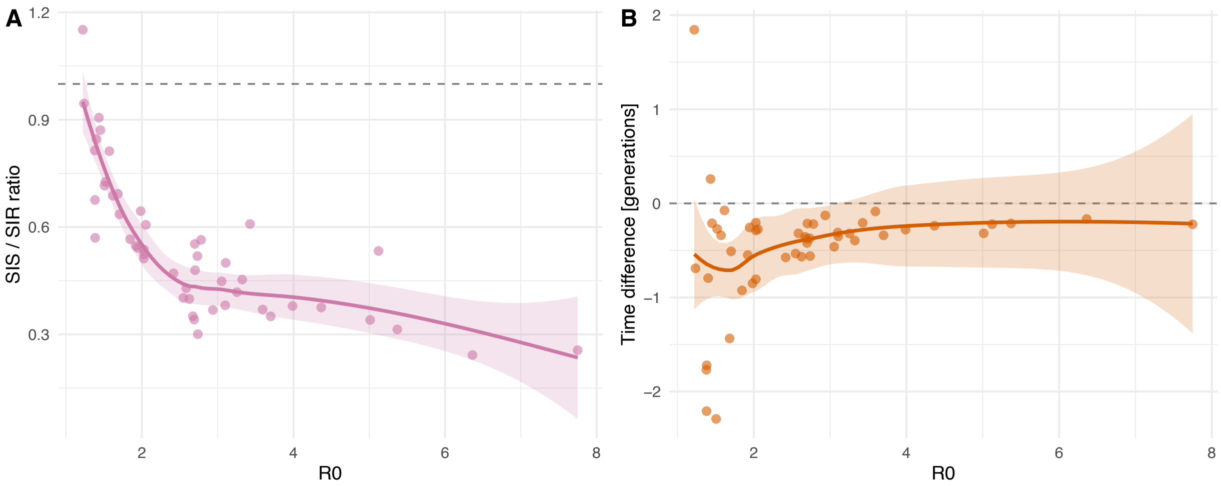


Figure S25: **Quantitative comparison of co-infection dynamics between SIR and SIS transmission models. A** Ratio of normalized co-infection rates between SIS and SIS models across different reproduction numbers ($R_0$). Each point represents a paired simulation, with the dashed horizontal line indicating equal co-infection rates (ratio = 1). Values above 1 indicate higher co-infection rates in SIS models, while values below 1 indicate higher rates in SIR models. The purple trend line with confidence interval shows the systematic relationship between $R_0$ and co-infection ratio differences. **B** Difference in peak co-infection timing between SIS and SIR models, expressed in generations (recovery rate × time difference). Positive values indicate that peak co-infection occurs later in SIS models compared to SIR models. The orange trend line reveals how transmission intensity ($R_0$) affects the temporal offset between peak co-infection timing in the two model types. These comparisons highlight fundamental differences in reassortment opportunity windows between populations with temporary immunity (SIR) versus populations with no acquired immunity (SIS).

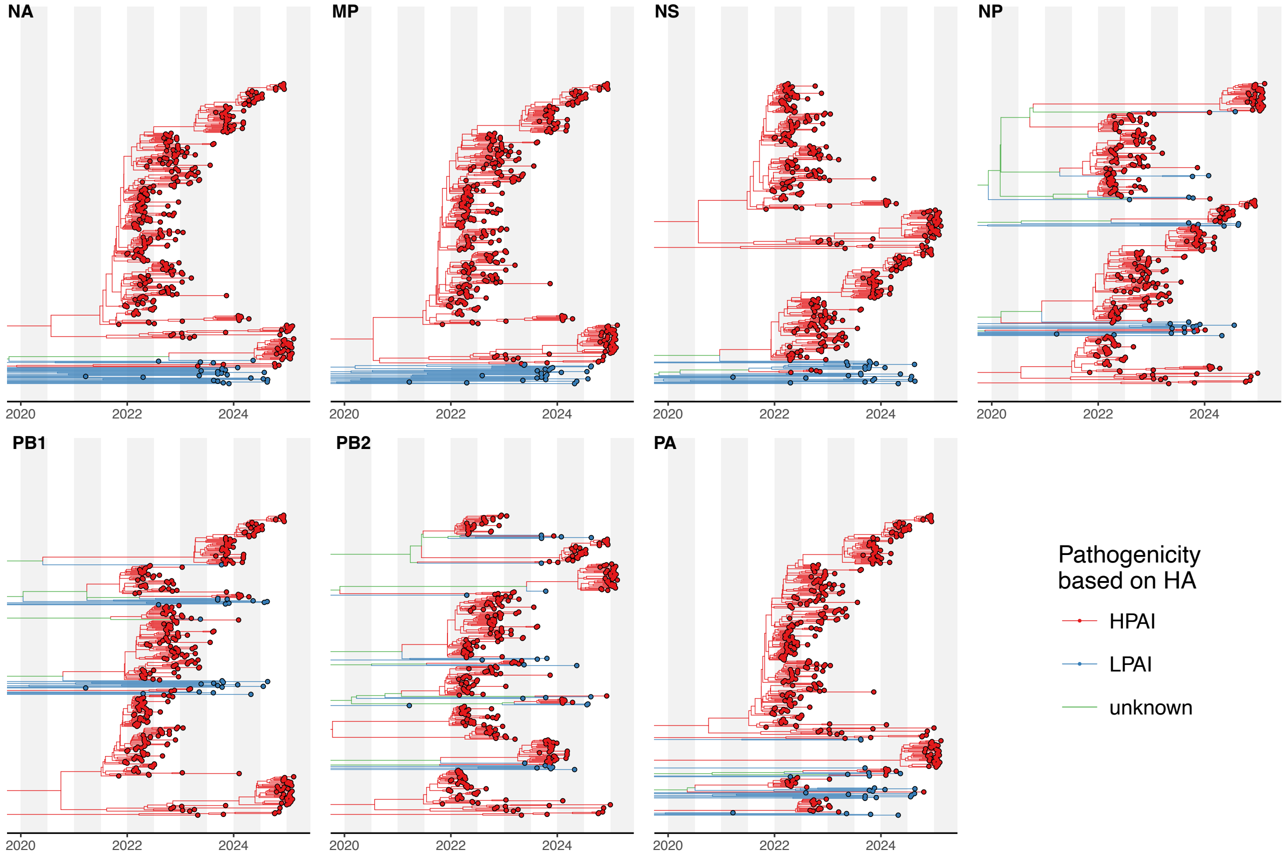


Figure S26: **MCC trees for non-HA segments of HPAI and LPAI H5N1.** Maximum Clade Credibility trees for the seven non-HA segments of HPAI and LPAI H5N1 sequences from North America. LPAI lineages are shown in blue (determined by the corresponding HA lineages), while HPAI lineages are shown in red.

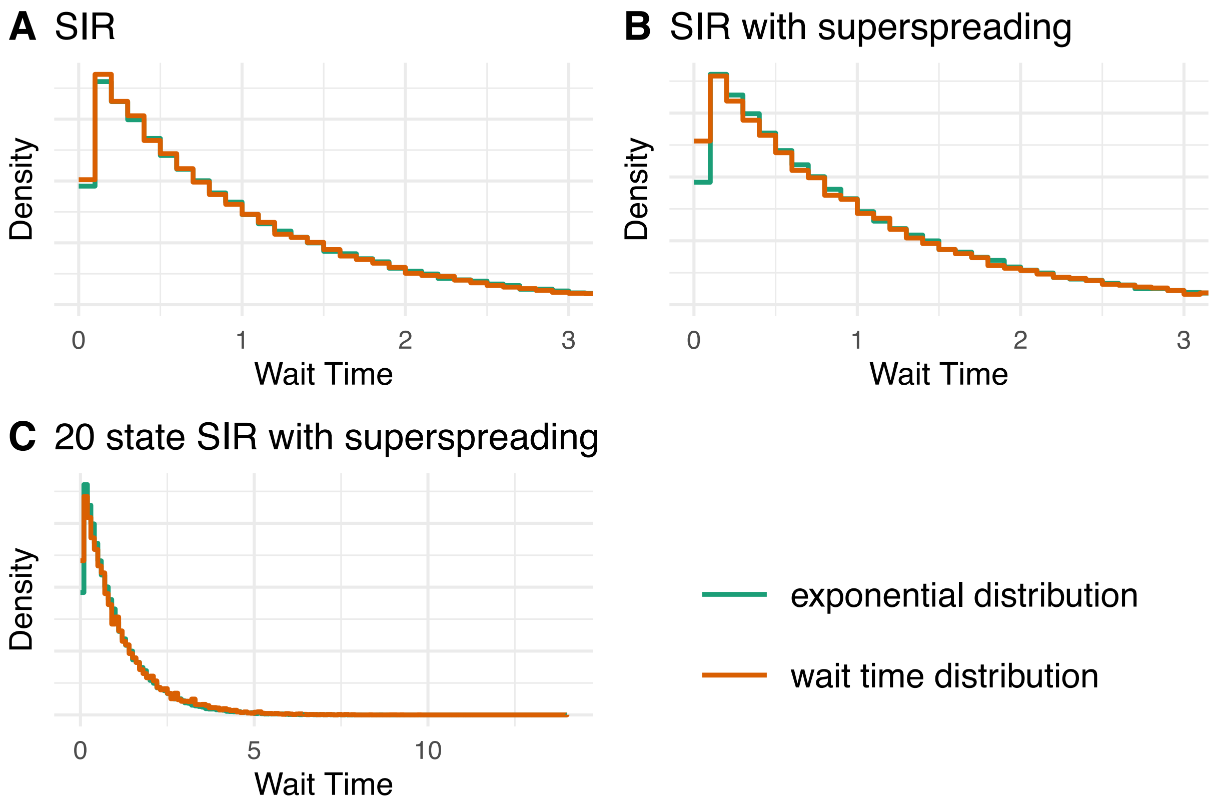


Figure S27: **Wait time distribution between reassortment events.** Here, we compare the distribution of wait times after correcting for disease prevalence and transmission rates using the expression for $\rho(t)$ to the probability density function of an exponential distribution with mean 1. The wait time distributions are calculated from reassortment networks simulated using **A** SIR models with co-infection, **B** SIR with co-infection and superspreading modelled using a Poisson distribution, and **C** a 20-state SIR model with spreading.

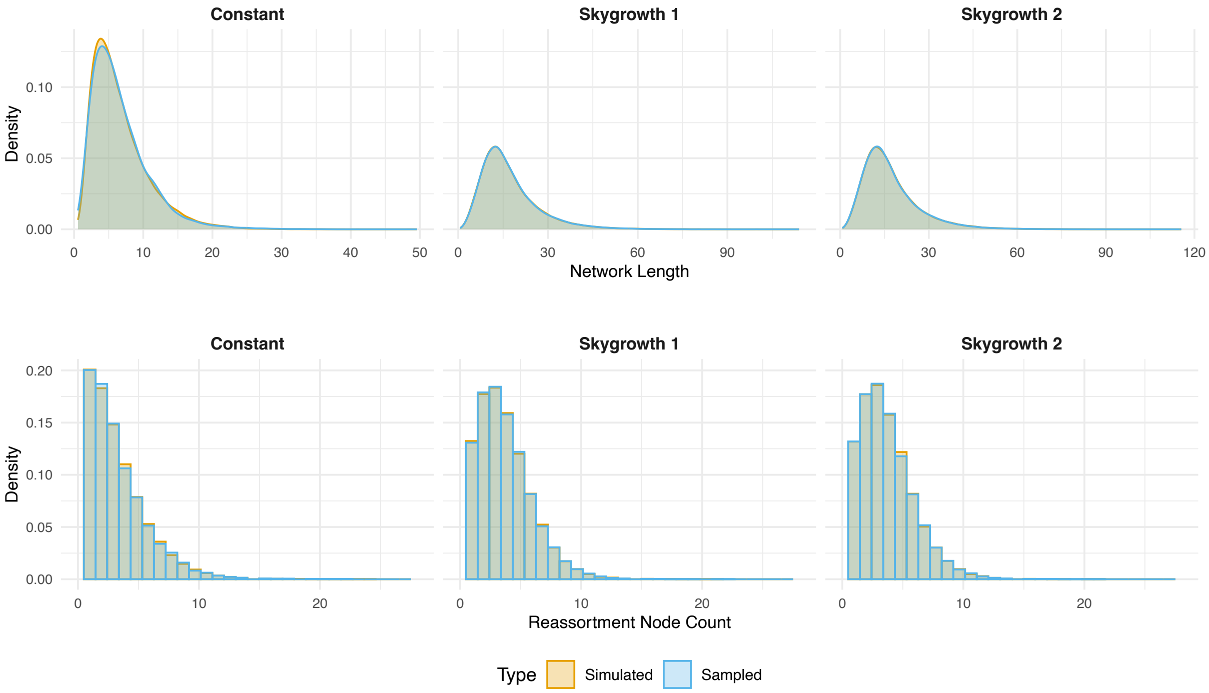


Figure S28: **Comparison of the network heuristics sampled under the prior distribution compared to simulated under the same distribution to validate the implementation of the Hastings ratios.** **A** Validation of the MCMC implementation by comparing summary statistics of reassortment networks sampled from the prior distribution (without data) against networks simulated directly under the same prior distribution. The agreement between these distributions confirms the correct implementation of the Hastings ratios and proposal mechanisms. Multiple network statistics are compared, including the number of reassortment events, tree balance measures, and temporal clustering, to ensure comprehensive validation of the inference framework.

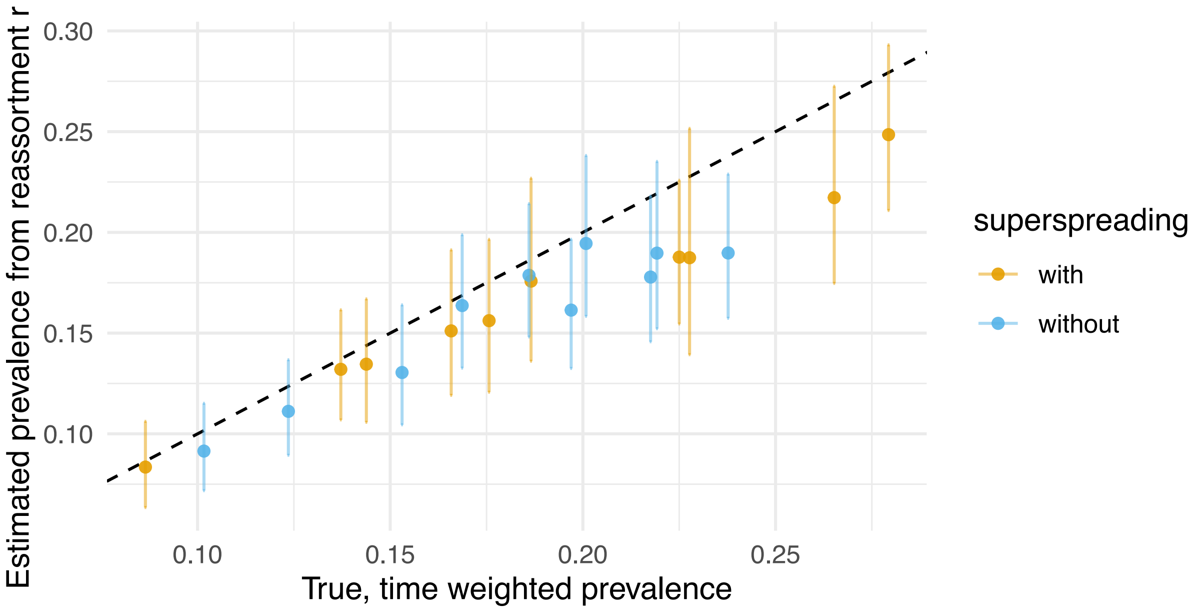


Figure S29: **Reassortment rate estimation from SIR models assuming constant reassortment rates over time. A** Comparison between true time-weighted prevalence (x-axis) and estimated prevalence from reassortment rate inference (y-axis) for simulations with (orange) and without (blue) superspreading. Points represent individual simulation runs, with error bars showing 95% credible intervals. The dashed line represents perfect agreement between true and estimated values. Results demonstrate the accuracy of constant rate estimation when the underlying transmission dynamics follow simple SIR models.

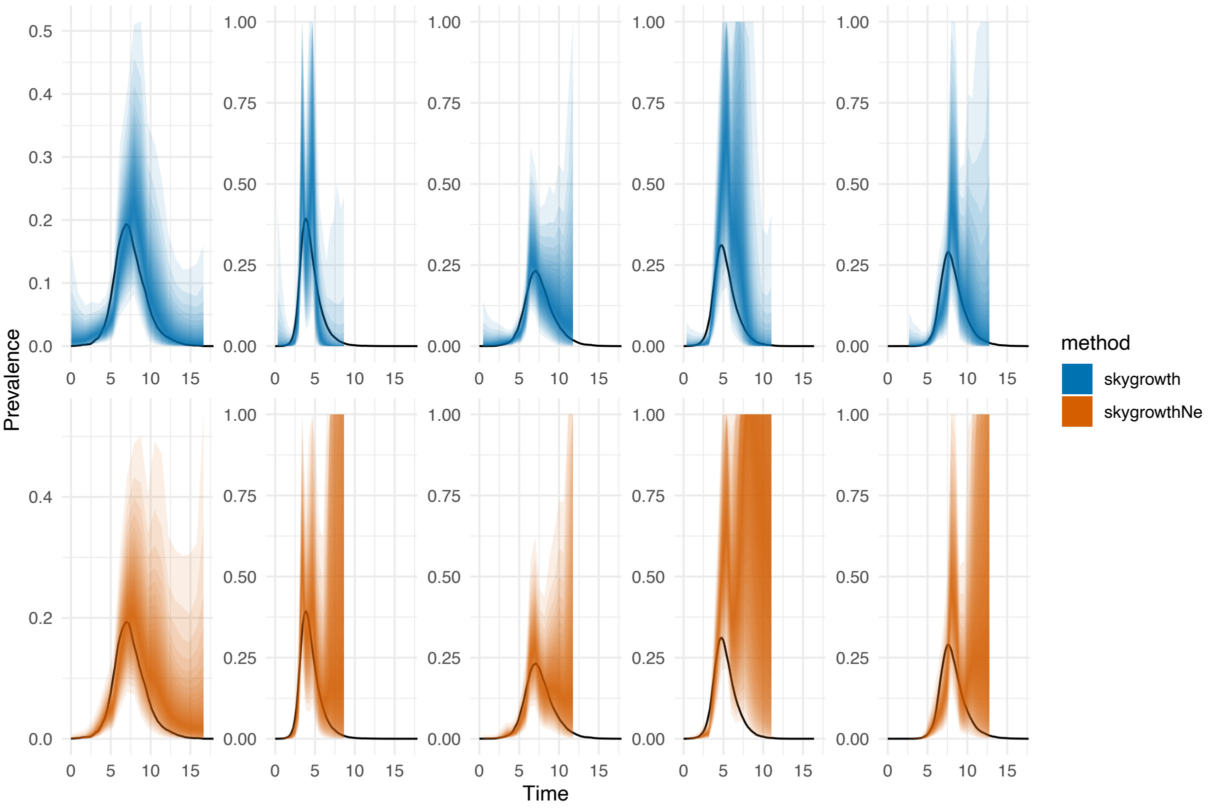


Figure S30: **Time-varying reassortment rate estimation from SIR models. A** Estimated time-varying reassortment rates (colored ribbons) compared to true prevalence trajectories (black lines) for 5 simulation runs without superspreading. Each panel represents a different simulation run, with blue and orange ribbons showing 95% credible intervals for two different estimation methods (skygrowth and skygrowthNe). The x-axis shows time relative to the most recent sampling time, and y-axis shows prevalence. Ribbons capture the true prevalence dynamics, validating the time-varying rate estimation approach.

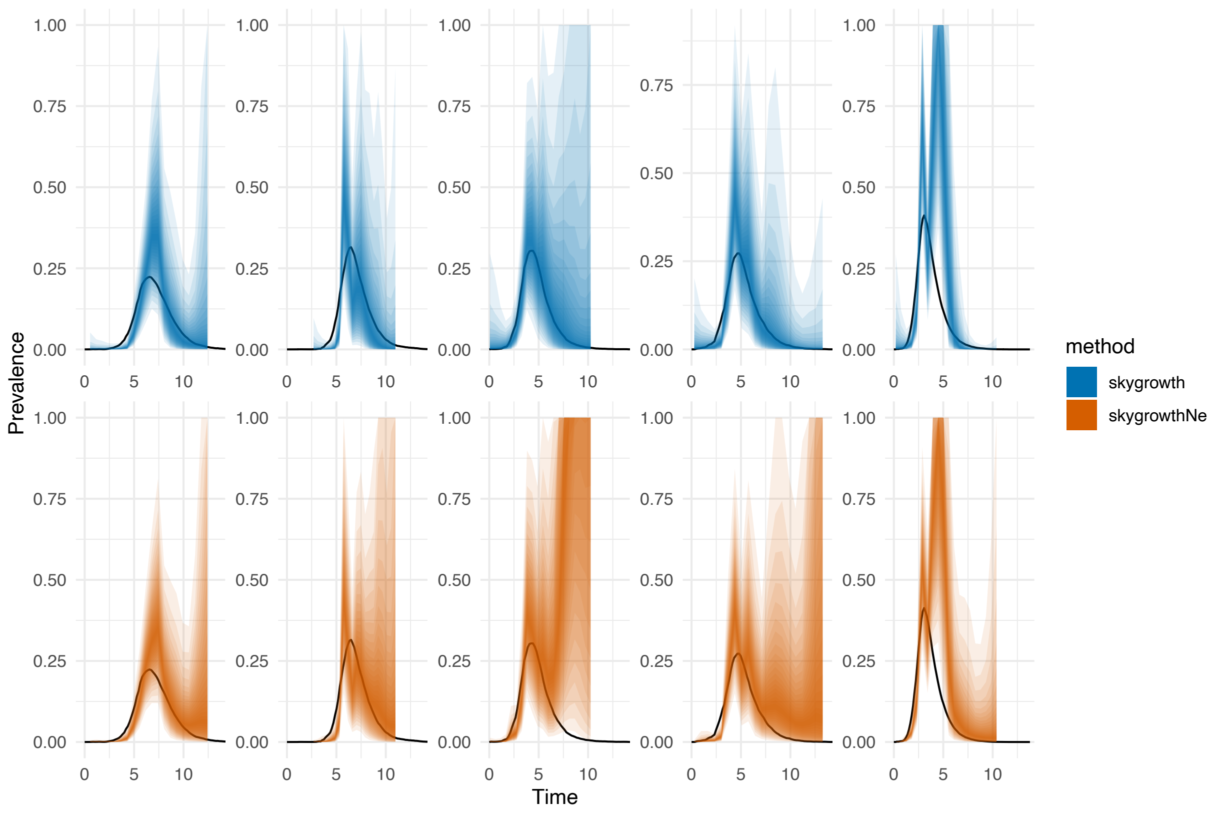


Figure S31: **Time-varying reassortment rate estimation from SIR with superspreading models. A** Estimated time-varying reassortment rates (colored ribbons) compared to true prevalence trajectories (black lines) for 5 simulation runs with superspreading dynamics. Each panel represents a different simulation run, with blue and orange ribbons showing 95% credible intervals for two different estimation methods. The presence of superspreading events creates more variable transmission dynamics, which are successfully captured by the time-varying rate estimation methods. Results demonstrate robustness of the approach even under heterogeneous transmission scenarios.

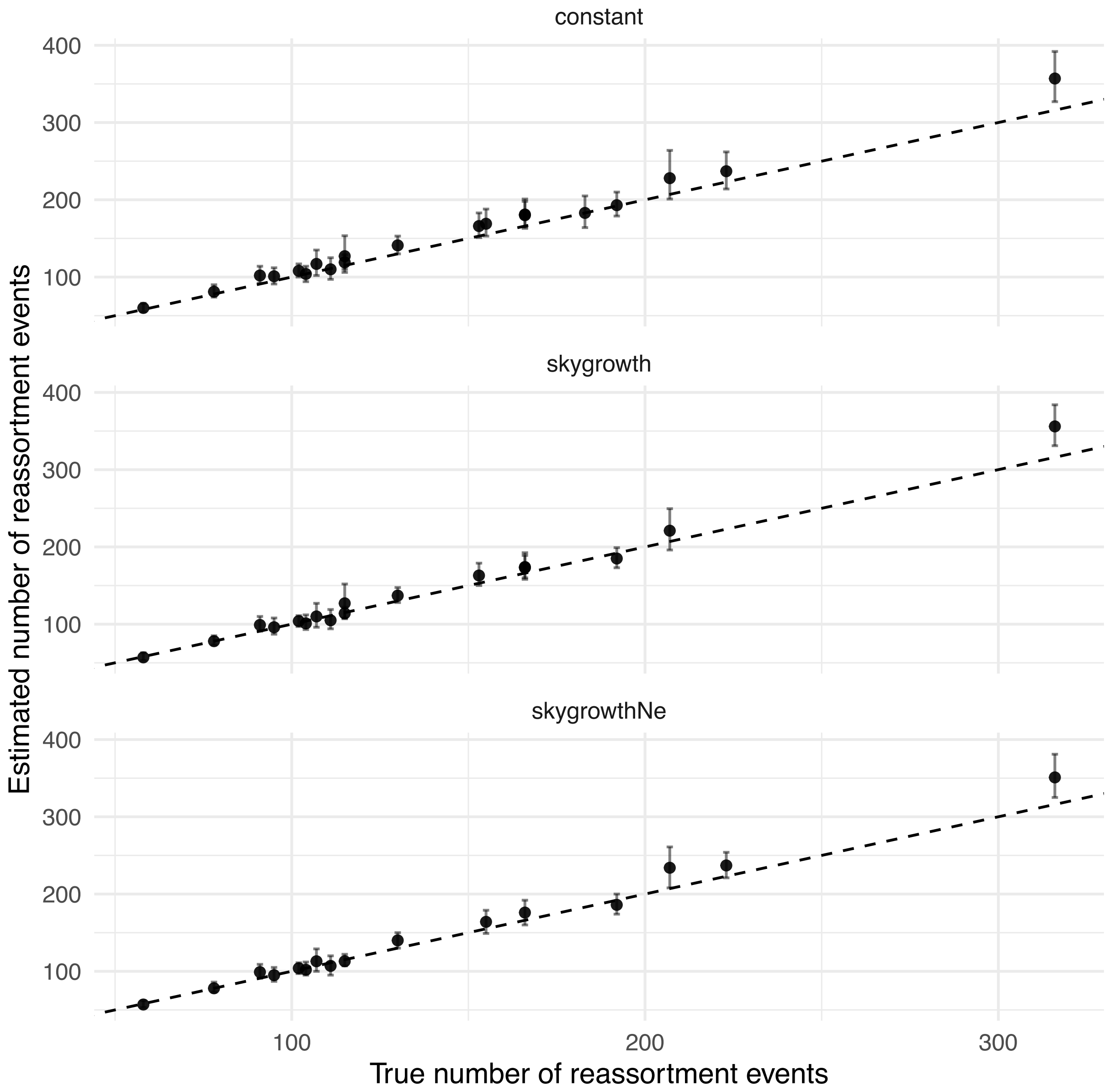


Figure S32: **Number of reassortment events estimated with and without accounting for time-varying reassortment rates and effective population sizes. A** Comparison of estimated versus true number of reassortment events across all simulation runs. Top panel shows results for constant rate estimation, bottom panel shows results for time-varying rate estimation methods. Points represent individual simulation runs with error bars showing 95% credible intervals. The dashed line represents perfect agreement. Both methods show good agreement with truth, with time-varying methods showing slightly improved accuracy for scenarios with dynamic transmission patterns.

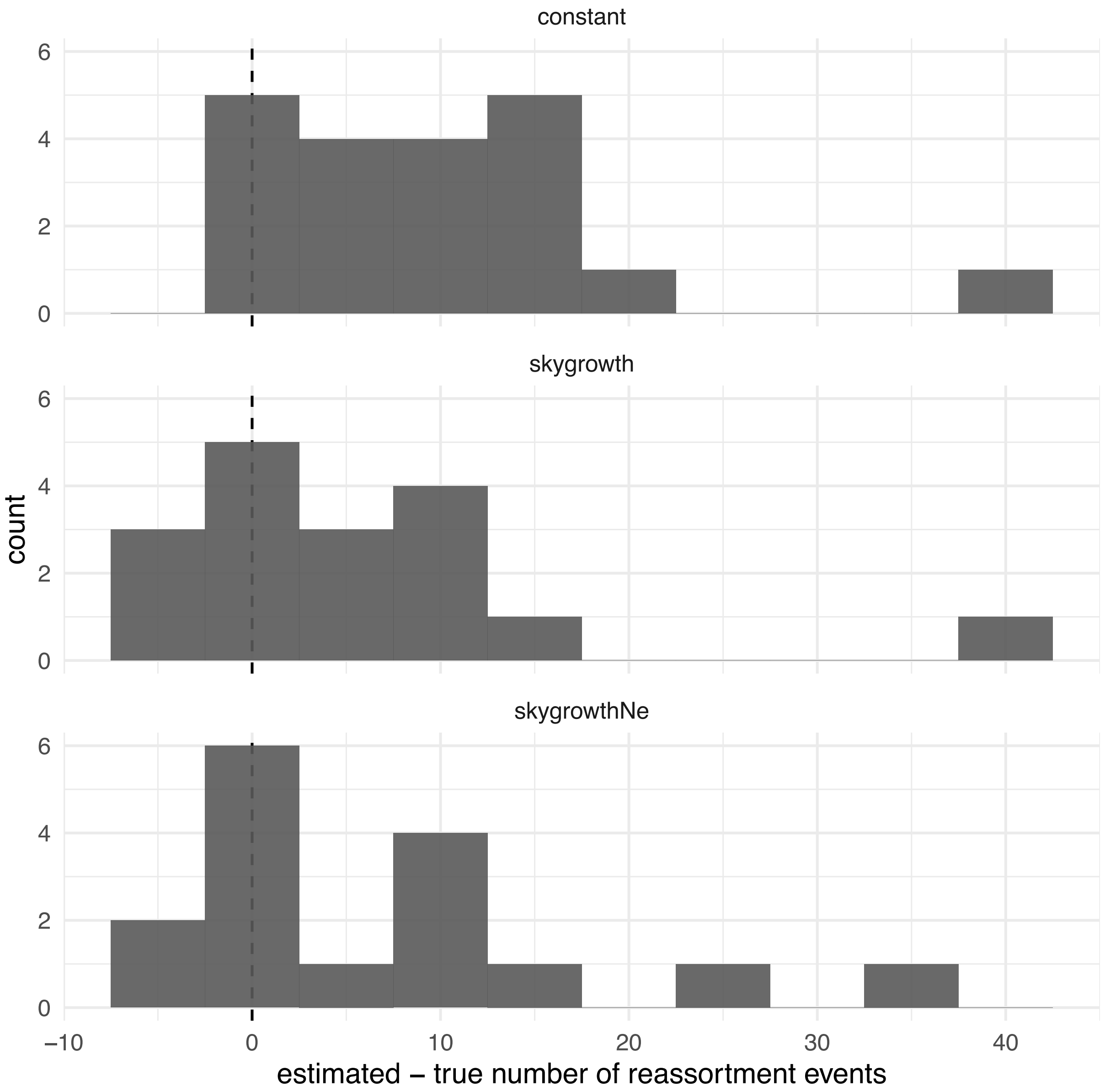


Figure S33: **Bias in the number of reassortment events estimated with and without accounting for time-varying reassortment rates and effective population sizes. A** Distribution of bias (estimated minus true number of reassortment events) for constant rate estimation (top) and time-varying rate estimation (bottom). Vertical dashed line at zero represents unbiased estimation. Histograms show the frequency of over- and under-estimation across simulation runs. Both methods show roughly unbiased estimation with the majority of runs clustering around zero bias, indicating that neither method systematically over- or under-estimates the number of reassortment events.

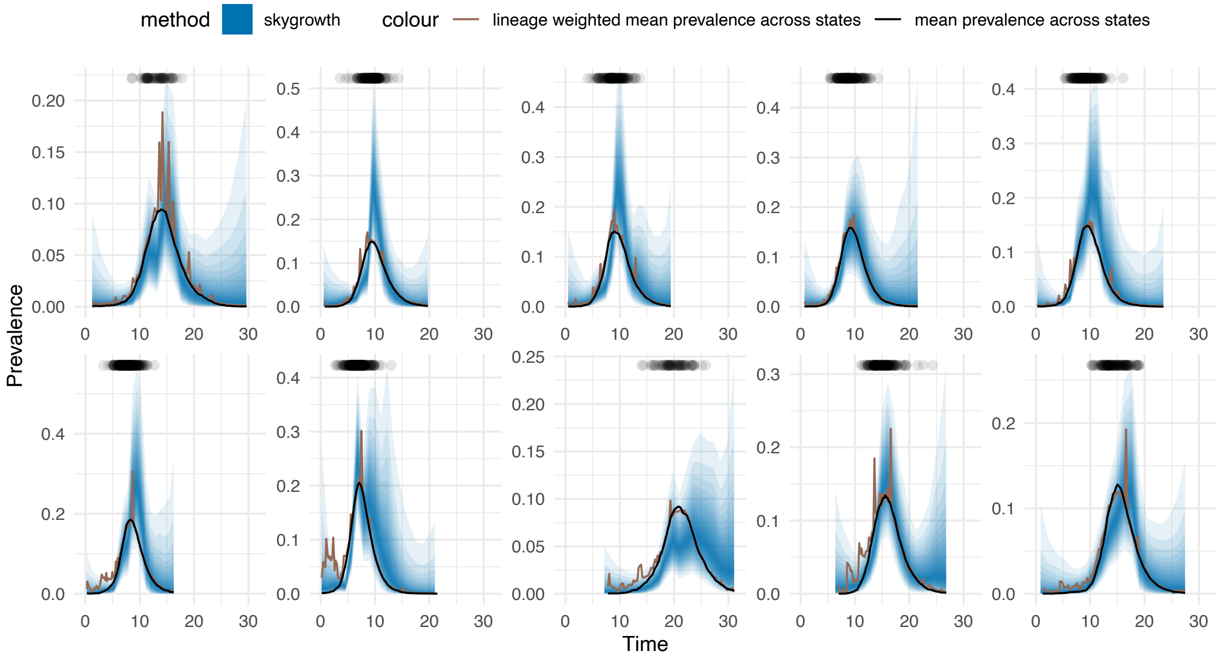


Figure S34: **Time-varying reassortment rate estimation from structured SIR models. A** Results from structured population models with multiple demes showing estimated time-varying reassortment rates (colored ribbons) compared to true prevalence trajectories. The structured models incorporate spatial or demographic heterogeneity in transmission, representing more realistic epidemiological scenarios. Multiple colored ribbons represent different quantile intervals of the posterior distribution, demonstrating uncertainty quantification in the presence of population structure.

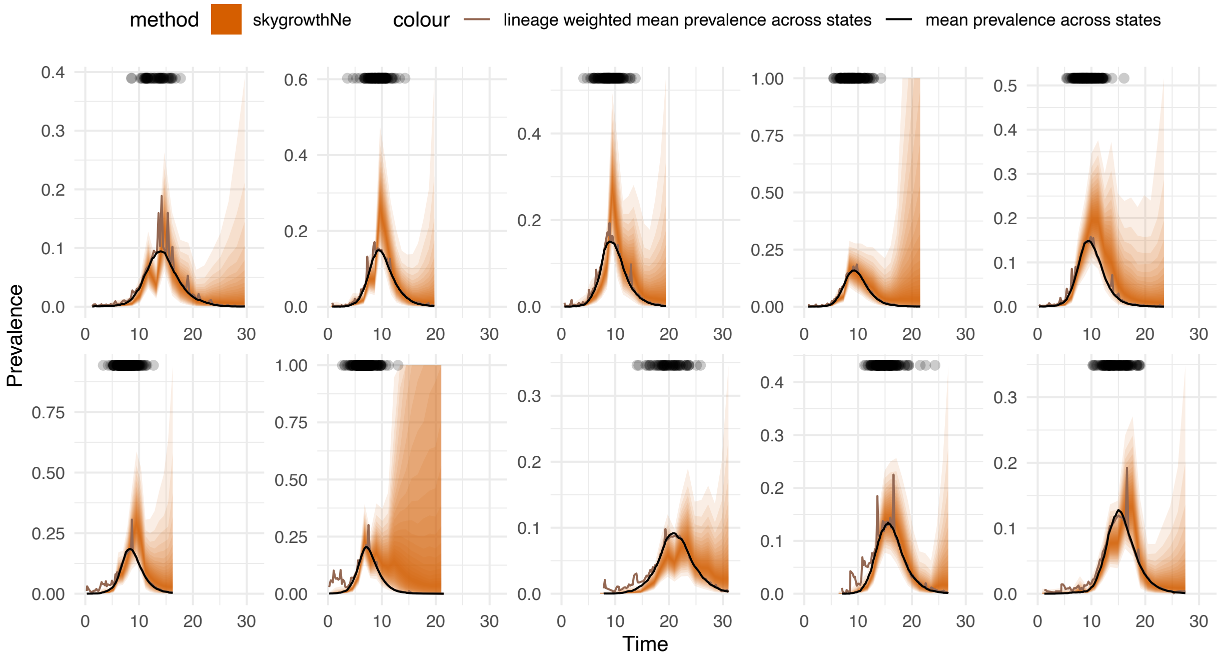


Figure S35: **Time-varying reassortment rate estimation from structured SIR models, when modeling the reassortment rates as a function of the effective population size. A** Results from structured SIR models where reassortment rates are explicitly modeled as functions of effective population size rather than simple prevalence. This approach accounts for demographic fluctuations that may not be captured by prevalence alone. Colored ribbons show credible intervals for the joint estimation of effective population size and reassortment rates, providing more nuanced inference about transmission dynamics in structured populations.

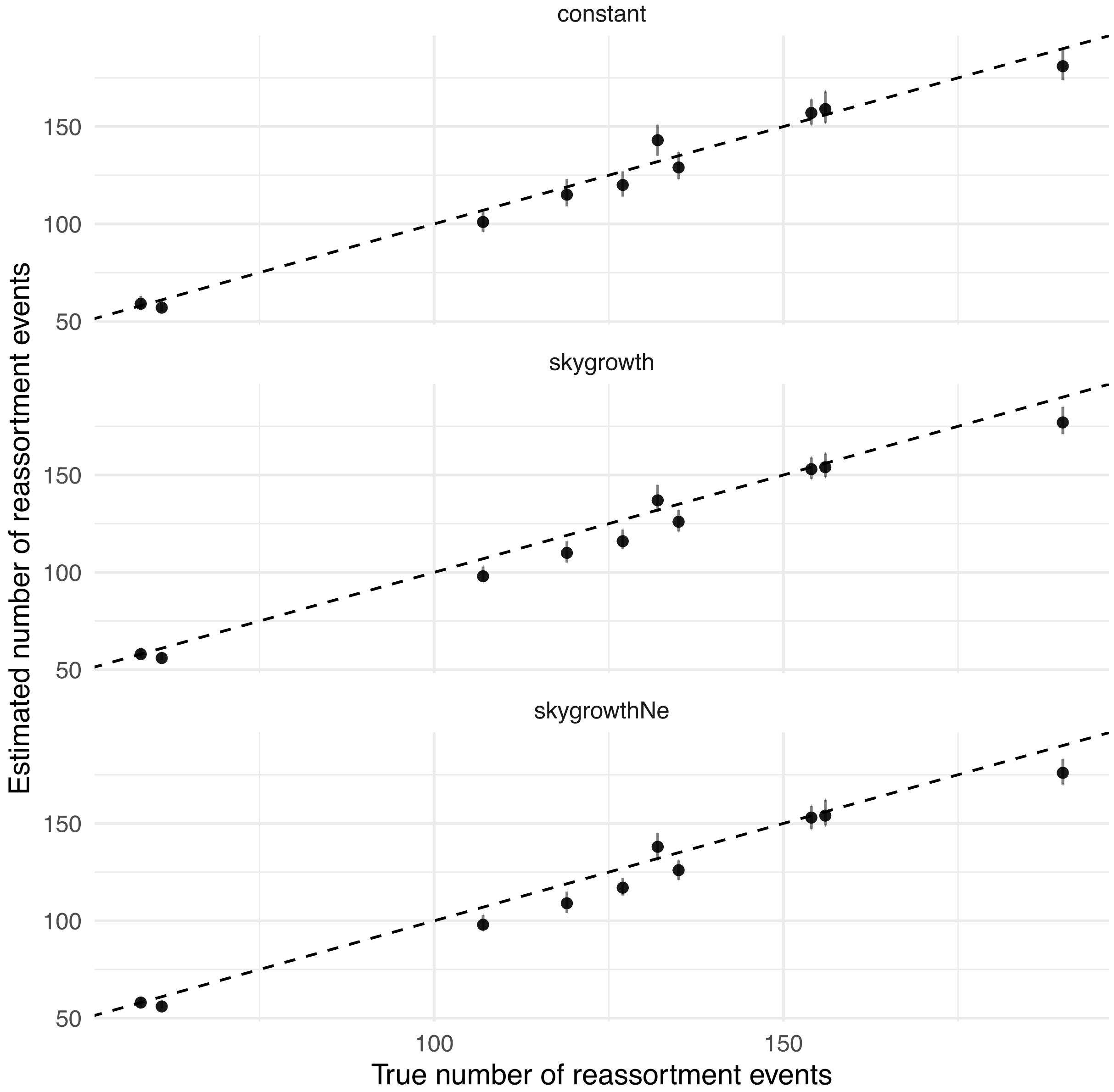


Figure S36: **Number of reassortment events estimated with and without accounting for time-varying reassortment rates and effective population sizes in structured populations. A** Validation of event count estimation in structured SIR models comparing estimated versus true numbers of reassortment events. Points represent individual simulation runs with error bars showing credible intervals. Results demonstrate that accounting for population structure and time-varying rates maintains accuracy in event count estimation even in complex demographic scenarios with multiple interacting populations.
